# Transposable elements strongly contribute to cell-specific and species-specific looping diversity in mammalian genomes

**DOI:** 10.1101/679217

**Authors:** Adam G Diehl, Ningxin Ouyang, Alan P Boyle

## Abstract

**Background:** Chromatin looping is exceedingly important to gene regulation and a host of other nuclear processes. Many recent insights into 3D chromatin structure across species and cell types have contributed to our understanding of the principles governing chromatin looping. However, 3D genome evolution and how it relates to Mendelian selection remain largely unexplored. CTCF, an insulator protein found at most loop anchors, has been described as the “master weaver” of mammalian genomes, and variations in CTCF occupancy are known to influence looping divergence. A large fraction of mammalian CTCF binding sites fall within transposable elements (TEs) but their contributions to looping variation are unknown. Here we investigated the effect of TE-driven CTCF binding site expansions on chromatin looping in human and mouse.

**Results:** TEs have broadly contributed to CTCF binding and loop boundary specification, primarily forming variable loops across species and cell types and contributing nearly 1/3 of species-specific and cell-specific loops.

**Conclusions:** Our results demonstrate that TE activity is a major source of looping variability across species and cell types. Thus, TE-mediated CTCF expansions explain a large fraction of population-level looping variation and may play a role in adaptive evolution.

## Background

Ever since chromosomes were first observed microscopically, it has been speculated that their 3D structure plays a central role in regulating nuclear function (1,2). Early observations revealed that individual chromosomes occupy distinct nuclear territories and, while their arrangement varies between different cell types, this structure is conserved between mother and daughter cells (2). These findings led to the hypothesis that chromosome structure directly influences cellular phenotypes. Since that time, microscopic and molecular studies have dissected chromatin structure into an intricate hierarchy of large-scale territories, compartments, domains, neighborhoods, and loops (3–8), confirming the importance of 3D structure in regulating gene expression, replication, and other nuclear processes. However, the mechanisms by which these structures are created and maintained, how they modulate gene expression, and how they evolve are still poorly understood.

A common feature of chromatin loops is the presence of insulator proteins at their boundaries, most notably CTCF (3,4,9–14). Although this property has been observed across distantly-related metazoan phyla (10), it is especially important in mammals, where CTCF knockdown leads to widespread loop disruption and gene dysregulation (15). There are several known cases of differential gene expression resulting from differential looping (16–21), Chromatin loops often direct enhancers to target genes and differential enhancer-promoter contacts can lead to differential gene expression (22). However, there is no consensus on the underlying mechanisms of this process and how they may be affected by changes in chromatin looping. Furthermore, while it is known that most mammalian loops are tethered by CTCF sites, how differential CTCF occupancy across cell types relates to loop specificity or how CTCF binding site turnover relates to interspecies looping divergence requires elucidation.

Comparisons between various human and mouse cell types have shown that differential looping is common (3,4,6), even between individual cells within a tissue (22). Differentially-looped regions are associated with changes in gene expression, coinciding with both cross-species and cross-cell expression variability between human, mouse, and chimpanzee (4,23,24) and aberrant expression associated with congenital diseases (25) and cancer (26). This suggests a mechanistic link between loop variation, gene expression changes, and adaptive evolution and there is evidence that CTCF binding site divergence contributes strongly to this variation (23).

A large fraction of binding sites for transcription factors (TFs), including CTCF, are derived from transposable elements (TEs) (27–30). TEs are mobile genetic elements that can influence observable phenotypes (31), often by altering the expression of nearby genes (32). Many TEs proliferate using a “copy-paste” mechanism, which allows them to undergo exponential amplification and disperse to new locations throughout their host genome. This creates DNA substrates upon which Darwinian selective forces can act to create evolutionary novelty. Recent studies have broadly implicated TE proliferation in gene regulatory innovation and the reasons for this appear to derive from innate features of TEs, including embedded TF binding sites (TFBSs) capable of influencing the expression of nearby genes (reviewed in (33)). There is evidence to suggest that cooption of embedded CTCF binding sites as loop anchors is a prominent mechanism in neofunctionalization of TE-derived sequences. TE-derived CTCF sites are frequently found at the boundaries of chromatin loops in mouse (3) and recent evidence has implicated TE proliferation in creating intraspecies variation in CTCF binding and cancer risk (34–37). However, the overall contribution of TE-derived CTCF sites to the evolution of genome folding across mammals and their relative contributions to conserved and divergent looping remain unknown.

In this study, we explore the extent to which TEs have affected CTCF binding and chromatin looping in the human and mouse genomes. We revisit the question of CTCF-binding enrichment in specific TE families and find that CTCF enrichment and neofunctionalization as loop anchors appear to be independent, with TE-enrichment correlated with the strength of the CTCF binding motif in the TE consensus sequence, while prevalence at loop anchors is a function of TE abundance. We further cataloged the shared chromatin loops in these cells with at least one anchor mapping to a TE-derived site from any TE family, demonstrating that an unexpectedly large fraction of loops are anchored within TEs. Importantly, whether loop anchors are within a recognizable TE or not, they are marked by similar patterns of conservation and activating histone marks characteristic of gene regulatory sequences and these marks are only found in cells with a detectable loop anchored at a given locus. We devised a system to classify loops by their breadth of use across cells and species and demonstrate that TE-derived CTCF sites predominantly associate with cell-specific and species-specific loops. We speculate that TE-driven CTCF binding site expansions throughout human and mouse evolution have contributed broadly to looping diversity and created novel enhancer-promoter contacts. The resulting increase in flexibility of gene regulatory programs may have allowed subpopulations to adapt to differential selective pressures, catalyzing divergence and specialization to distinct evolutionary niches.

## Results

Our ultimate goal was to investigate the impact of transposable element (TE) proliferation on 3D chromatin structural divergence. To do so, we identified chromatin loops in the human and mouse genomes in which at least one anchor was derived from a transposable element. By classifying loops by their degree of sharing between species and cell types we determined the contribution of TEs to cross-cell and cross-species looping divergence. We further investigated the properties and potential effects of TE-driven loop divergence by identifying sets of loops in which a species-specific TE insertion created a differential loop. An example of such a region is shown in Figure 1. We chose to focus on loops anchored by CTCF sites because of its well-characterized role in chromatin looping (3,5,11,14,15,19,38–45) and known enrichments within several families of SINE, LINE, and LTR retrotransposons in multiple mammalian species (27–29).

**Figure 1:**
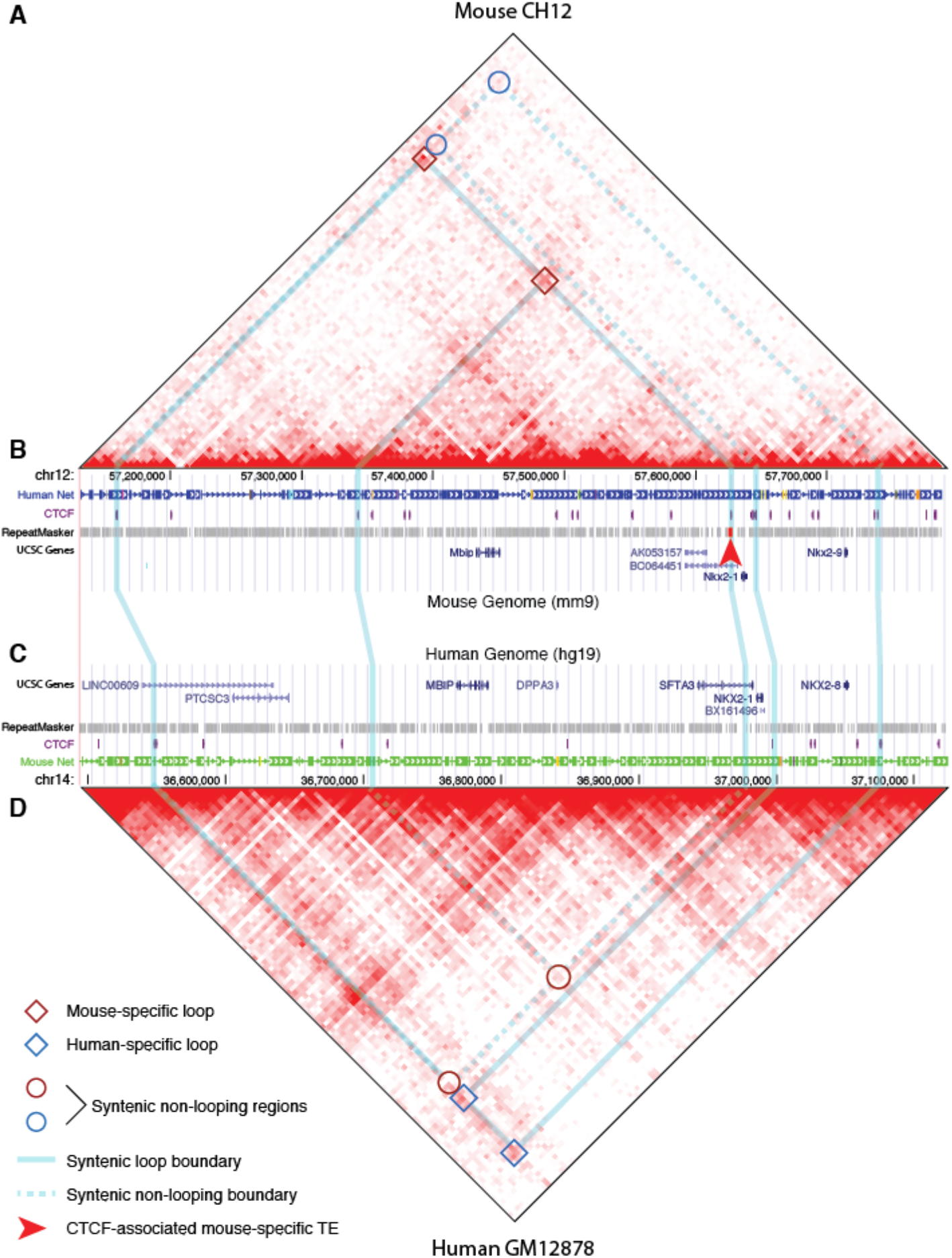
Transposable element insertions create novel, species-specific loop contacts. Hi-C maps for a differentially-looped region in human and mouse are shown in panels A and D. X axes indicate genomic positions along a syntenic region of mouse chromosome 12 and human chromosome 14. 3D loop contacts are visible as clusters of bright red pixels away from the X axis while bright red areas along the X axis mostly represent experimental noise. **(A)** Mouse CH12 Hi-C map showing two mouse-specific loops (dark red boxes). The syntenic locations of human-specific loops are indicated by blue circles. **(B)** UCSC Genome Browser track for mouse, showing relevant features within this genomic region. Both mouse loops are tethered by CTCF binding sites, with the shared right anchor falling within a mouse-specific retrotransposon (bright red bar marked by arrowhead). **(C)** UCSC Genome Browser track for the orthologous region in the human genome. Syntenic locations of loop anchors in the mouse and human genomes are connected by vertical blue lines. **(D)** The human GM12878 Hi-C map for this region shows the absence of looping between the conserved left anchor and the region syntenic to the mouse-specific right anchor (dark red circles). Two human-specific alternative loops are indicated by blue boxes.

### Human and mouse CTCF binding landscapes are strongly influenced by TE activity

We first assessed the genome-wide effects of TE proliferation on CTCF binding in human and mouse. CTCF ChIP-seq data for matched immune cell types from both species (table S1) were combined into a union set of orthologous and species-specific binding sites and intersected with known TE insertions (46) (Figure 2A). The results show that TEs have contributed strongly to CTCF binding in both species, constituting ~35% of all CTCF binding sites. CTCF binding was highly variable across species, with >85% of sites showing species-specific occupancy and TEs contributed a surprisingly large fraction of these sites. Overall, TE-derived CTCF sites comprise >47% of mouse-specific sites (>36 times more than expected by chance) and >30% of human-specific CTCF binding sites (>82 times more than expected by chance). TE-derived sites appear to show a preference for species-specific occupancy, regardless of whether the insertion occurred before or after speciation, suggesting a strong link between TE activity and CTCF binding divergence.

**Figure 2:**
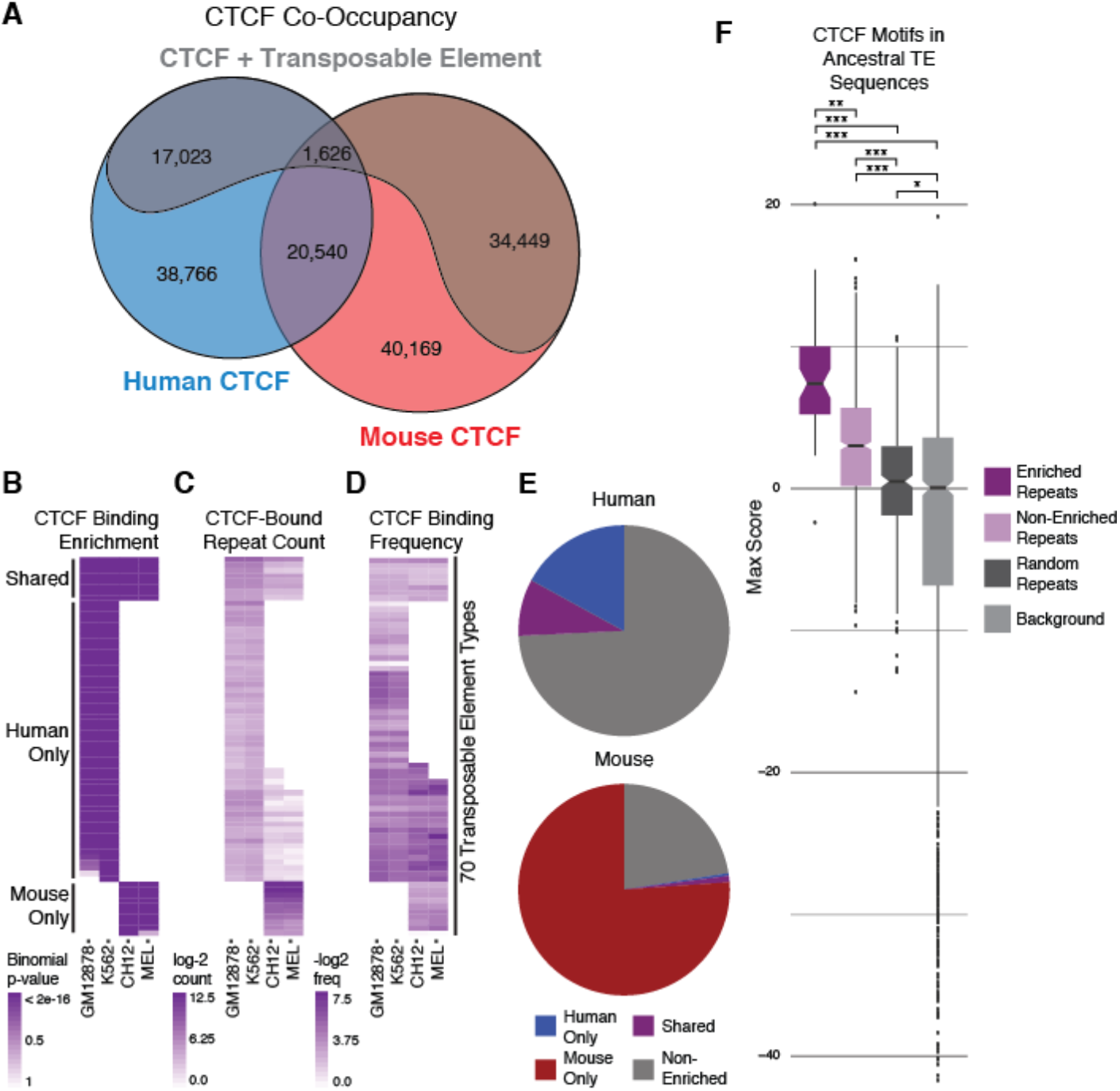
CTCF binding variability is associated with transposable element activity. **(A)** Proportion of CTCF sites in the human and mouse genomes with conserved and divergent binding and their respective transposable element (TE)-derived fractions. Human-specific and mouse-specific fractions include both orthologous and non-orthologous CTCF binding sites. **(B)** Binomial tests recovered 70 TE types significantly enriched for CTCF binding in at least one cell type. Enrichments were classified as human-only, mouse-only, or shared based on the cell types in which they were observed. **(C)** Counts of CTCF-bound copies of each enriched TE type per cell type. **(D)** Percentage of detectable copies bound by CTCF for each enriched TE type in each cell type. **(E)** Fractions of TE-derived CTCF binding sites originating from human-enriched, mouse-enriched, shared, and non-enriched TE types in human and mouse. **(F)** Log-odds score distributions for the strongest CTCF motif match within consensus sequences (a proxy for the ancestral TE sequence) for CTCF-enriched and non-enriched TEs, compared to randomly-selected TEs and length-matched background sequences. Scores above 1 represent sequences with greater than random resemblance to the CTCF motif. Wilcoxon rank-sum tests were used to test for significant score differences in all pairwise combinations: *** p = 2.2e-16, ** p = 1.6-14, * p = 0.03.

Previous studies have found strong CTCF-binding enrichments within several families of SINE, LINE, and LTR retrotransposons in multiple mammalian species (27–29). In our analysis, we noted that several previously unreported subfamilies from these TE classes (we call these TE types) had made substantial contributions to CTCF binding in both human and mouse. Given this observation, we wanted to revisit the question of CTCF binding enrichment within our dataset. We adapted enrichment testing methods used in three previous studies (27–29) and compared the results. Overall, we observed good agreement between results from all three methods (Figure S1, Tables S3, S4). Because the binomial testing method produced the most conservative results, we chose to use these data for all subsequent analyses.

Binomial testing yielded 70 CTCF-enriched TE types (Figure 2B, Table S2). Despite being performed on different cell types than those used in previous studies, our tests captured over 76% of CTCF-enriched TE types reported previously, showing that CTCF binding enrichments are highly robust and reproducible. Enrichment strength was correlated with neither TE abundance (Figure 2C) nor CTCF binding frequency (Figure 2D), and all but a few enrichments spanned both cell types within a species, showing that CTCF-binding enrichments are not cell-type specific (Figure S2C-D). Our methods recovered 60 previously unobserved TE types, likely due to the larger size of the present dataset compared to previous studies. All TE types in which we observed enrichments were previously identified in recent a genome-wide screen for regulatory exaptation of TE elements in human (47). However, whereas that study identified exaptation events from all known families of TEs, we saw only a subset of these in our data: L1 LINEs; Deu and B2 SINEs; ERV and Gypsy LTRs; hAT and tcMar DNA elements, and two types of mouse-specific L1-dependent retrotransposons (Table S2).

While these observations agreed well with previous reports, we also observed that a large fraction of both human and mouse CTCF binding sites fall within instances of TEs from families not enriched for CTCF binding (Figure 2E). In fact, all but two of the major classes (LINE_Merlin and DNA_Dong-R4) found by Haussler and Lowe (47) were represented in our dataset, even though only a subset of these were CTCF-enriched. Given the robustness of our tests, we are confident that non-enriched types do not represent false negatives, leading us to speculate that only presence of a latent CTCF motif within a TE is required for exaptation as a binding site. Consistent with this hypothesis, we observed that recognizable CTCF motifs are present in the consensus sequences (a proxy for the ancestral TE sequence) for both enriched and non-enriched TE types (Figure 2F). In fact, 38% of TE consensus sequences in Repbase (48) contain a latent CTCF motif, suggesting that CTCF may play a yet unknown role in replication and/or transposition of these elements.

Notably, while the motif scores found in consensus sequences for non-enriched TE types were uniformly lower compared to enriched types, this was the only significant difference we could find between the two classes. We saw no significant trends in a comparison of CTCF binding strength, motif scores, and phastCons conservation scores for across extant instances of both classes of TEs (Figure S2). This suggests that the only functional difference between enriched types relative to their non-enriched counterparts is that their ancestral forms may have bound CTCF more strongly, perhaps allowing them to more readily divert binding from nearby existing binding sites. Nevertheless, motifs within extant copies of enriched TE types did not score higher, on average, than those belonging to non-enriched types (Figure S2B), suggesting that binding affinity within both classes is pushed by selection towards an optimal level. We speculate that, while strong CTCF binding may be required for a TE type to show statistical enrichment for CTCF binding, the chance of exaptation for function is likely dependent on other factors, such as location, local chromatin environment, and proximity to genes, regulatory elements, and existing CTCF binding sites.

Surprisingly, not all TE types enriched for CTCF binding only in human (human-only types) arose after divergence from mouse. In fact, the major dispersals for nearly half of human-only types are ancestral, i.e., predating human-mouse divergence, in some cases by over 100 million years (Figure 3A). We considered two hypotheses to explain this anomaly: 1) that additional primate-specific dispersals of these TE types may have occurred after divergence from rodents, or 2) that differential selective pressure on insertions present in the most-recent common ancestor (MRCA) led to progressive loss of CTCF binding in mouse. To investigate the first hypothesis, TE insertion ages were estimated for all enriched TE types using previously published methods (27) (Figure 3A). As expected, age distributions for most human-only and mouse-only TE types were centered near or below the estimated primate-rodent divergence date. By contrast, all but four ancestral TE types enriched in human-only had median dispersal dates predating the divergence date by at least 25 million years. Furthermore, the human and mouse age distributions for most of these were similar, suggesting that they originated from the same dispersals. Only six types showed age distributions consistent with subsequent human-specific dispersal.

**Figure 3:**
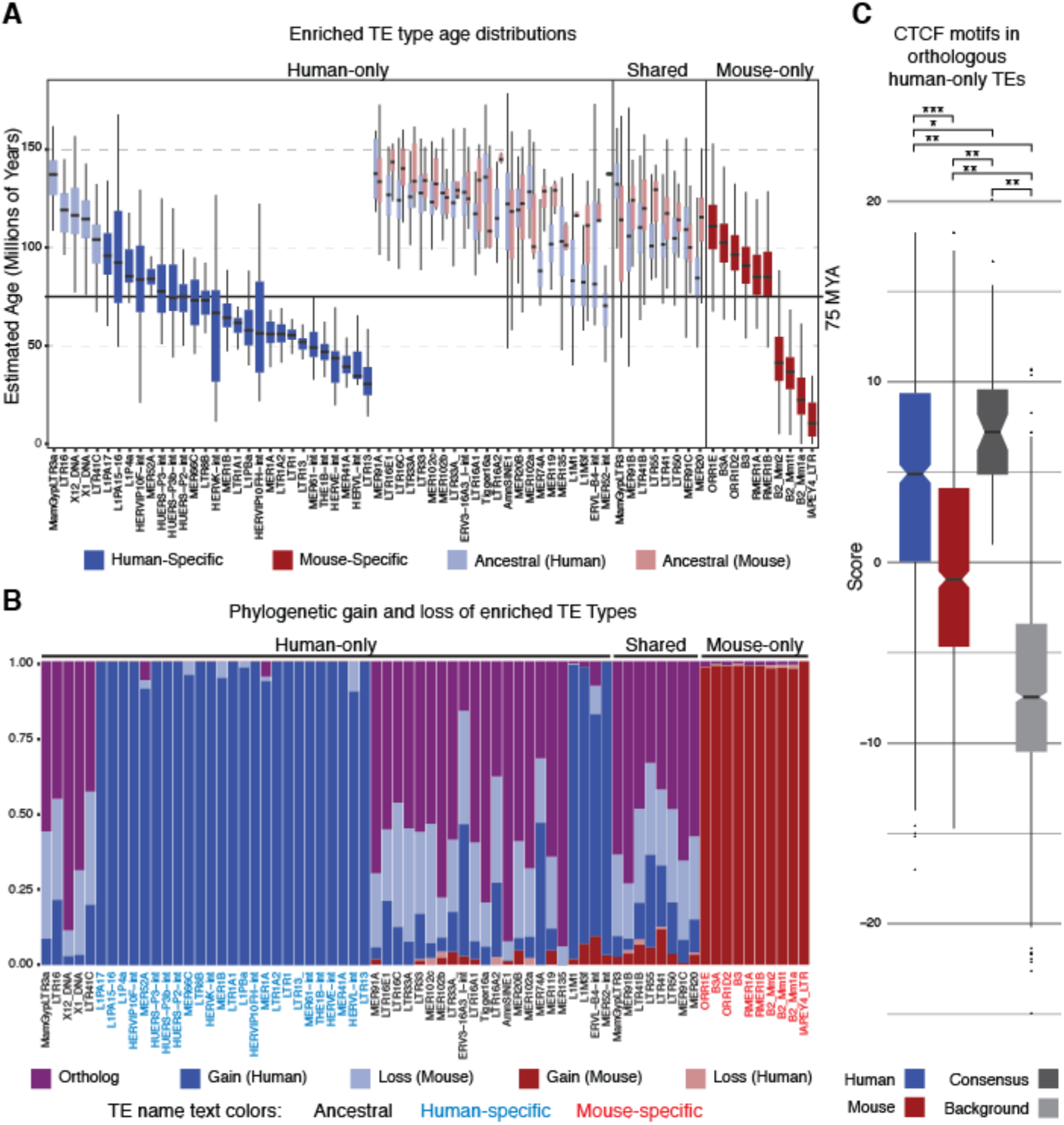
Ages and phylogenetic histories of human-only CTCF-enriched TEs support mouse-specific loss-of-function. **(A)** Estimated age distributions for CTCF-bound TE insertions of enriched types. Colors indicate the species-specificity of each TE type and score distributions are split by species where applicable. The solid black horizontal line marks the estimated primate-rodent divergence date. **(B)** Fraction of enriched TE insertions inferred as orthologous, or as evolutionary gains or losses on a given branch of the phylogeny. TE label colors indicate whether the given type is ancestral (black), human-specific (blue), or mouse-specific (red). **(C)** Maximum CTCF motif scores within human and mouse instances of ancestral, human-only TE types. TE Consensus sequences (a proxy for the ancestral TE sequence, dark gray) and length-matched background sequences (light grey) are shown for comparison. Scores greater than 1 indicate sequences matching the CTCF motif more than would be expected by chance. Hypothesis tests were used to assess significance of differences in all pairwise comparisons: *** p = 2.9e-58 (Wilcoxon signed-rank test), ** p < 4.9-19, * p = 2.6e-4 (Wilcoxon rank-sum tests).

While extant copies of human-only ancestral TE types are detectable in the mouse genome (49), they are bound by CTCF much less frequently than in human (Figure 2C). In fact, five of these TE types showed no evidence for CTCF binding within any mouse copies. Because most human and mouse insertions appear to have originated from the same dispersals, it seems likely that differential selection has led to loss of CTCF binding within mouse copies while human copies retained functionality. To further investigate this possibility, we performed a phylogenetic gain/loss analysis to determine the fraction of ancestral human-only TE type insertions present in the MRCA (Figure 3B). Consistent with our hypothesis, orthologs and sequences lost from the mouse genome after speciation constitute the overwhelming majority of these loci. Given this result, we wanted to know if these loci retain the ability to bind CTCF in mouse. The maximum CTCF motif scores within human and mouse sequences for all extant orthologous instances of these TE types were compared. Consistent with loss of function, mouse instances scored systematically lower than their human counterparts and >2/3 had maximum motif scores below the theoretical threshold for CTCF binding. By contrast, the majority of human instances had maximal scores that suggested robust CTCF binding ability (Figure 3C). While the consensus sequences for all these TEs contain strong CTCF motifs, these data suggest that mouse instances have progressively lost the ability to bind CTCF after divergence from the MRCA, explaining the observed differential enrichments. It is possible that subsequent massive dispersals of mouse-specific B2 and B3 SINEs, which have no parallel in human evolution, have driven this loss of function by diverting CTCF binding from nearby ancestral sites, resulting in relaxed constraint.

### TE-activity has strongly contributed to chromatin looping in human and mouse

We next wanted to determine the contribution of TE-borne CTCF sites to human and mouse loop anchors. RAD21 ChIA-pet loops from human GM12878 and K562 cells and Hi-C loops from mouse CH12 cells (Table S1) were filtered for CTCF presence and trimmed to coincide with the strongest embedded CTCF ChIP-seq peak at each anchor, then intersected with TE annotations (46). This allowed us to determine that 28% of loops across all types were derived from TE-borne CTCF sites (Figure 4A), and nearly 10% of these are formed entirely from TE-derived anchors (Figure 4B). We observed that, consistent with previous reports (3,7,44), ~88-89% of loops are formed between pairs of convergent or inward-pointing CTCF motifs, regardless of cell type or TE content (Figure S3, Table S6). Importantly, both TE-derived and non-TE-derived loops exhibit similar CTCF-binding properties (Figure 4A-C) and patterns of activating histone marks (Figure S4). Consistent with our hypothesis that CTCF-enrichment does not directly influence exaptation as a loop anchor, we found that the fractions of enriched and non-enriched TE types contributing to human and mouse chromatin loops are proportional to their overall contributions to CTCF binding (Figure 4D). Indeed, all but nine of the TE families present in our CTCF dataset have contributed chromatin loop anchors in human and/or mouse, with their overall contributions roughly scaling with their abundance. This strengthens our conclusion that gain of function requires only a latent CTCF binding site, and any TE meeting this requirement can influence looping and contribute to regulatory divergence.

**Figure 4:**
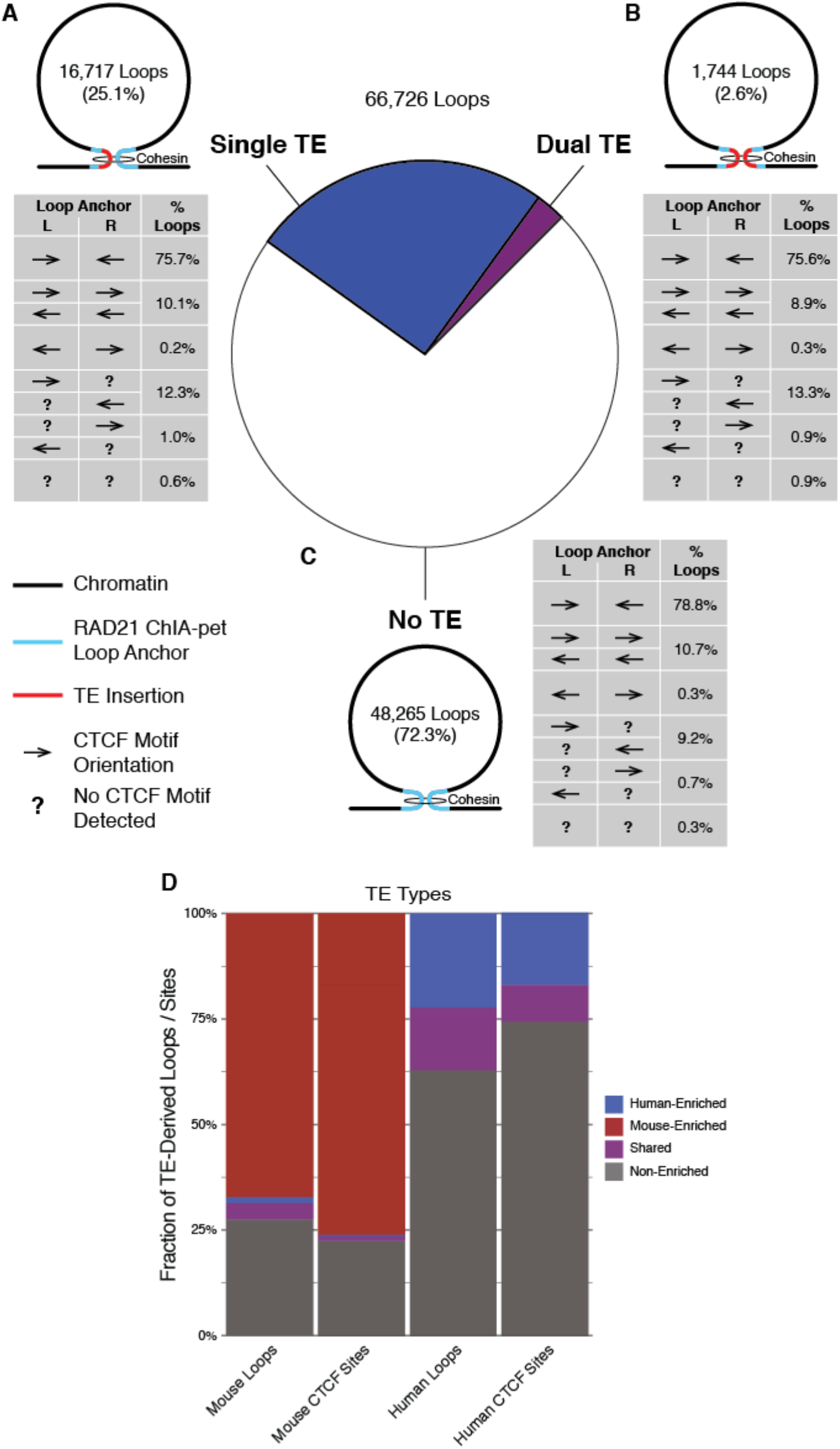
Transposable elements contribute strongly to chromatin looping in human and mouse. Human RAD21 ChIA-pet and mouse Hi-C loops containing CTCF ChIP-seq peaks at both anchors are broken down according to the number of TE-derived anchors they include. Each possible configuration is shown as a pictograph, with associated counts, alongside the central pie chart showing the fraction of all loops contributed by each configuration. Tables show the prevalence of different CTCF motif arrangements for each loop configuration. **(A)** Prevalence and CTCF motif arrangements for loops formed between pairs of TE-derived and non-TE anchors. **(B)** Prevalence and CTCF motif arrangements for loops formed between pairs of TE-derived anchors. **(C)** Prevalence and CTCF motif arrangements for loops formed between anchors not derived from known TE insertions. **(D)** Contributions of CTCF-enriched and non-enriched TE types to TE-derived loops and CTCF binding sites in human and mouse.

### CTCF-mediated chromatin loops are highly variable across cells and species

To further explore the relationship between TE-borne CTCF sites and structural evolution, we designed an algorithm to classify loops based on their degree of conservation across cells and species (Figure S5). Seven discrete conservation classes were defined based on cross-species mappability and overlap between annotated loop anchors in pairs of query and target cells (Figure. 5A). Loops within all pairs of query and target cells were compared with this method, allowing us to describe the conservation of CTCF-mediated looping in much greater detail than previously possible. Our results show that chromatin looping is highly variable across species and cell types. Less than 25% of loops were fully conserved between human and mouse and <50% of loops were fully conserved between GM12878 and K562 cells (Figure 5B). These rates are substantially lower than previous reports of 55-84% loop conservation across human cells, and 45-76% between human and mouse (3,4). However, it should be noted that previous methodologies employed a more relaxed definition of conservation than ours, requiring only that both query loop anchors map to *any* pair of anchors in the target cell, regardless of whether that pair forms a coherent, fully conserved target loop. Approximating this definition by aggregating data from classes C, B2, and B1 from our results yields conservation estimates very close to those reported previously (4).

**Figure 5:**
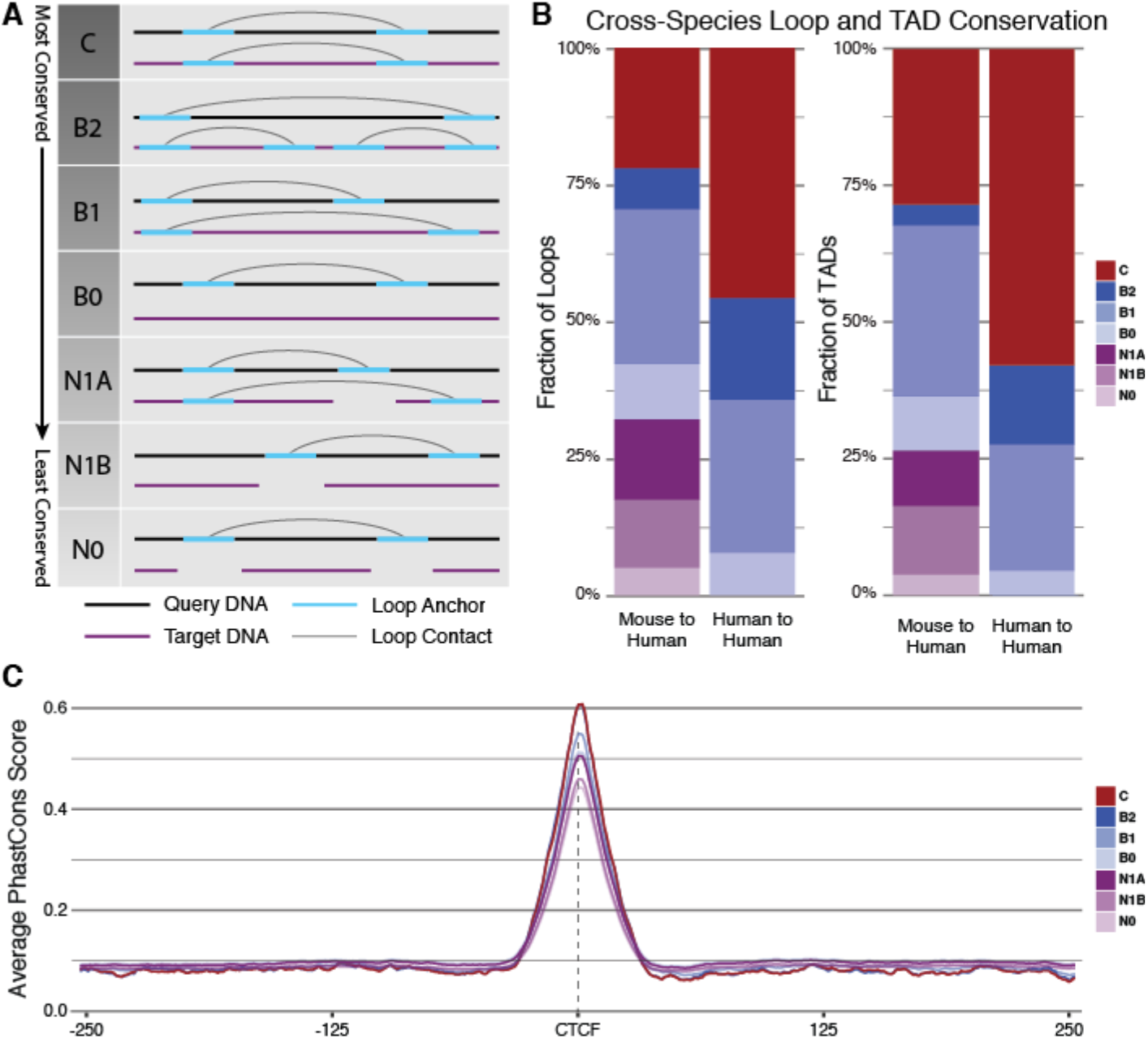
Conservation classes describe loop co-ocurrence patterns across cells and species, and are correlated with underlying phylogenetic conservation. **(A)** Conservation classes describe varying degrees of loop conservation between a query cell and a target cell based on sequence mappability and presence of a loop anchor at the syntenic locus. Diagrams illustrate the possible arrangements when comparing a query loop to a syntenic region in the target cell, with their corresponding conservation class labels. **(B)** Conservation class assignments from all pairwise comparisons between different cell types were aggregated into human-human and mouse-human comparisons. Plots show the contributions of each conservation class to the complete dataset (left), and to the subset of loops corresponding to known TADs (right). **(C)** Average phastCons conservation scores in 500bp windows centered at the CTCF ChIP-seq peak summit within human GM12878 loop anchors plotted for each conservation class, with CH12 used as the target cell.

Conservation classes also reflect underlying phylogenetic conservation within loop anchors, visible as a well-defined peak of PhastCons scores centered at the CTCF binding site which declines in magnitude with decreasing loop conservation (Figure 5C). This same pattern remained evident when only TE-derived loops were considered (Figure S7). This pattern may be partially explained by the younger evolutionary age of many species-specific loop anchors and decreased constraint on divergent ancestral elements. However, it is surprising that even the least conserved classes of loops show such strong evidence for functional constraint.

Previous studies have reported that strong conservation across cells and species is a hallmark of topologically-associating domains (TADs) (3,4). With this in mind, we separated known TADs from the rest of our dataset and plotted their conservation classes (Figure 5B, right panel). Surprisingly, conserved classes were only modestly enriched among TADs and, while the ~60% conservation we observed between human cell types is in line with previous reports (4), conservation across species was only ~30% -- roughly half the level of human-mouse conservation reported in the same study (4). However, their methods only required overlap between individual domain boundaries, not complete domains, and using their definition yielded comparable results for conservation across species (Figure S6). These results are consistent with the recent hypothesis that TADs are dynamic structures with a high degree of variability between species, cell types, and individual cells (8,24).

### TE-derived CTCF-sites are a primary source of looping variability

To further investigate the role of transposons in loop divergence, loop conservation data were combined with TE data and their relative contributions to each conservation class was calculated. While TEs contribute to loops in all conservation classes, their abundance increases as conservation decreases (Figure 6A-B). Given that our dataset includes many species-specific TE insertions, this result was not unexpected in the mouse-human comparison. However, the same trend was also evident between human cell types (Figure 6B, Figure S8B). This suggests that TEs are preferentially coopted into cell-specific functional roles, independent of when TE insertions occurred, which is in keeping with a recent study that identified TE enrichments at TAD boundaries that were dynamic over the course of cell differentiation (50). Importantly, we saw no significant differences in age distributions of TEs in any of the four conservation classes in comparisons between human cells (Figure 6E), suggesting that cooption into cell-specific roles arises as a direct consequence of falling within a TE. This appears to be true regardless of whether or not a TE descends from a CTCF-enriched family, as the relative contributions of enriched and non-enriched TE types to each human-human conservation classes are roughly proportional to their overall contributions to CTCF binding (Figure 6C).

**Figure 6:**
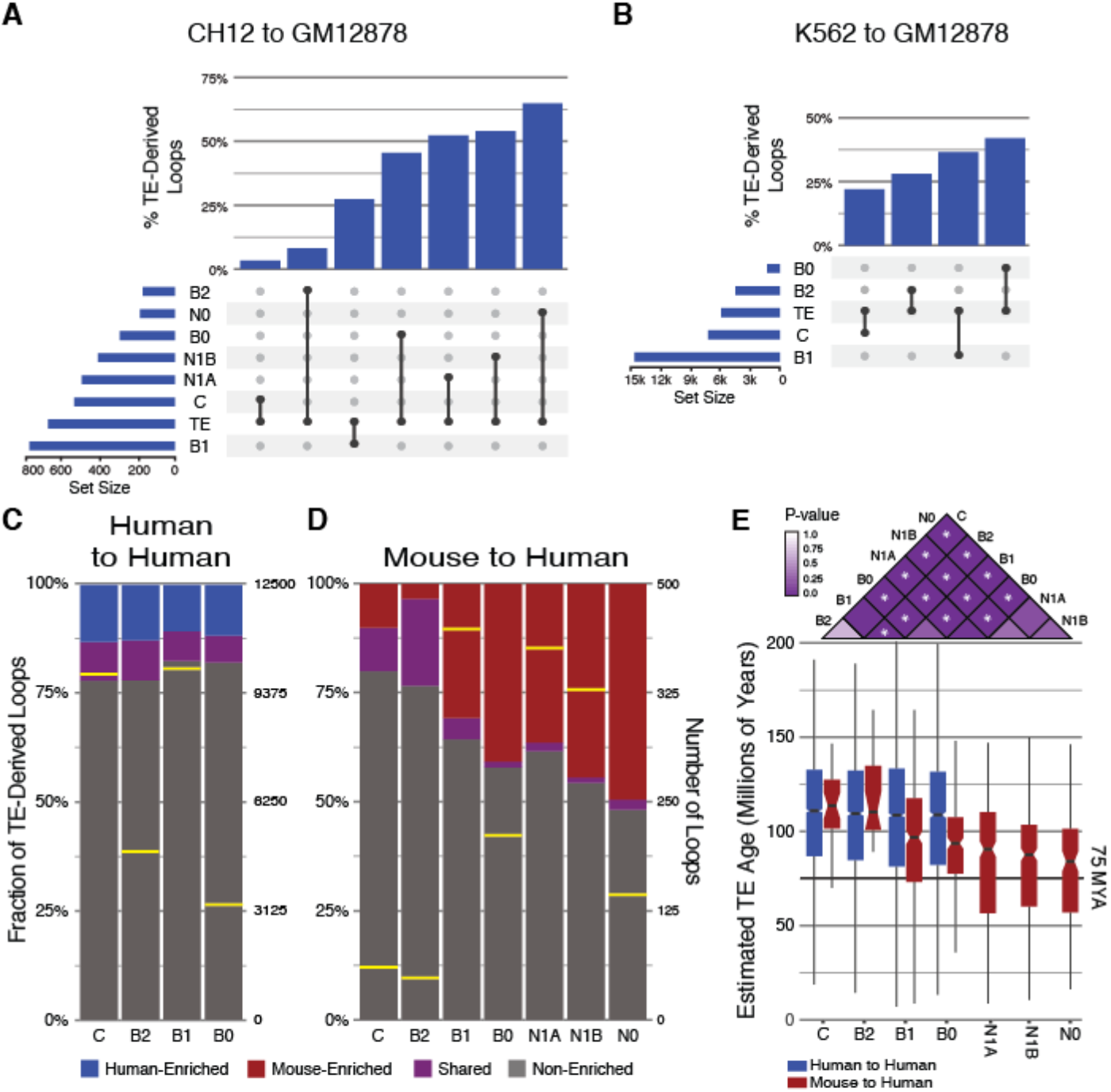
Transposable elements are correlated with species-specific and cell-specific loop formation. **(A)** UpSet plot showing TE-derived fractions within each conservation class for the comparison of mouse CH12 query loops to human GM12878 target loops. Horizontal bars show the number of loops assigned to each conservation class and the observed number of TEs among query loops. Vertical bars show the fraction of loops derived from TEs within each conservation class, ordered from left to right by decreasing structural conservation. **(B)** Same as (A), but between human K562 and GM12878 cells. **(C)** Fraction of TE-derived loops contributed by CTCF-enriched TE types in each conservation class for aggregate data from human-human comparisons. Yellow bars indicate the number of loops observed in each conservation class (right scale). **(D)** Same as (C), but for aggregated data from mouse-human comparisons. **(E)** Age distributions of TE insertions found at loop anchors in each conservation class for mouse-human (red) and human-human (blue) comparisons. The heat map shows Wilcoxon rank-sum test p-values for all pairwise comparisons between mouse-human conservation classes. P-values <= 0.01 are indicated by stars.

By contrast, CTCF-enriched TE types were markedly enriched in mouse-specific loops, especially those with anchors not mapping to human (Figure 6D). This likely reflects recent widespread dispersals of B2-class SINE elements in mouse. Consistent with this hypothesis, we observed a trend toward younger estimated TE ages in non-conserved loops relative to conserved loops in the mouse-human comparison, with significant differences present between most pairs of score distributions (Figure 6E). Notably, the age distributions of N1A, N1B, and N0 classes in mouse did not differ significantly from one another, mirroring the distributions seen in the human-human comparison and suggesting that the same set of TE dispersals contributed most of the loops in these classes.

Finally, we wanted to determine if cell-specific loops were associated with corresponding cell-specific functional signatures. All available histone modification data (Table S1) were compiled for overlapping loop anchors from GM12878 and K562 loops in the B0 class and compared to length-matched background sequences. If TE-derived cell-specific loops are associated with cell-specific functions, we would expect to see differential presence of activating histone marks in cells with a loop compared to those without. Indeed, GM12878-specific loop anchors had markedly stronger signal for several activating histone marks when compared to K562, in which no loops were detected between these loci (Figure S9). Thus we conclude that structural divergence introduced by TE-derived loops is also associated with functional divergence.

## Discussion

We present evidence that transposable elements have broadly impacted 3D genomic structure and contribute strongly to divergent chromatin looping in human and mouse. It appears that TE dispersals have shaped the 3D genomic landscape throughout mammalian evolution by distributing CTCF binding sites to novel locations in the genome. These sites may influence 3D genome structure by diverting CTCF from existing binding sites, thus creating novel loop contacts that in turn alter gene regulation. Although most studies to date have focused on the impact of CTCF-enriched TE types to the genome, we find evidence that both enriched and non-enriched TE types are likely to affect chromatin looping in proportion to their abundance. This appears especially true for human, in which CTCF-enriched TE types represent a small fraction of TE-derived CTCF binding sites. This has led to the apparently widespread misconception that TE activity has had relatively little effect on CTCF binding within primates. Our finding that ~1/3 of all CTCF-bound sites can be definitively attributed to a TE insertion combined with our observation that estimated age distributions for the TEs involved extend back well beyond rodent-primate divergence show that TEs have, in fact, broadly affected CTCF binding throughout primate evolution. Nevertheless, enriched TE types have had a particularly strong effect on the mouse looping landscape, likely owing to their sheer quantity. It seems probable that widespread dispersal of a TE that contains a strong CTCF consensus motif will affect genome-wide CTCF binding equally as broadly by diverting CTCF protein from nearby existing sites leading to overrepresentation among TE-derived CTCF sites. However, while such dispersals have broadly affected mouse and potentially other species (27,28,51), our results show human-specific CTCF enrichments to span a much larger number of TE types, each having undergone relatively smaller expansions. Why this is the case and whether it is also true of other mammalian clades are matters for future investigation.

We were surprised by the breadth of TE classes and families contributing functional CTCF binding and loop anchoring sites, which led us to speculate about the possible relevance of these sites to the TEs themselves. It seems unlikely that these sites would exist within the TE consensus sequences absent a function to the TEs themselves. It seems possible that these TE elements may harness the ability of CTCF to form loops in order to facilitate their own replication and/or integration into their host genome. For instance, these elements may rely on looping in order to bring them in contact with the host’s transcriptional machinery. Alternatively, they may utilize the association between CTCF and double strand breaks (52) to facilitate integration into the host genome. Regardless of the mechanism, these elements appear to exist in a symbiotic relationship with their hosts, harnessing host transcriptional machinery to proliferate while contributing substrates for regulatory innovation.

While TADs were originally described as stable structures with high conservation across species and cell types (3,4), our findings are consistent with newer reports showing TADs to be more dynamic. For example, single-cell assays have shown substantial variability between TAD boundaries among individual cells of the same type (22,53). The authors proposed that TAD formation is a stochastic process and experimentally observed TAD boundaries represent the average configuration over many cells. Even early Hi-C results found evidence for looping variability across subsets of the cell population (3) and this hypothesis may explain those findings. Furthermore, computer simulations have shown that chromatin loop extrusion, currently the most prevalent proposed mechanism for loop formation, requires that loops are dynamic and variable in order to recapitulate observed chromatin contact maps (45,54,55). Our results show that individual loop anchors rarely interact with only one downstream partner, which is consistent with the mechanism suggested by these studies.

Throughout this study we saw a strong connection between TEs and variability. Most importantly, TE-derived anchors were more likely to contribute to variable loops across species and cells. This finding coincides with other recent reports of associations between TEs and dynamic TAD boundaries (50). It is unclear what it is about TE-derived loop anchors that predisposes them to forming dynamic and variable loops, but this ability appears to facilitate their cooption as key regulators of metabolic and developmental genes (51,57) It is possible these TE-derived loops facilitate adaptive responses by increasing the number of alternative chromatin conformations around these genes. Our results are consistent with this hypothesis and suggest that transposable elements have contributed variability to the looping landscape at key points in mammalian evolution (Figure 3A). Indeed, it has been observed that divergence dates frequently coincide with retrotransposon dispersals, often with relatively greater TE insertions on one branch following speciation (56,57). Looping variability introduced by novel TE-derived CTCF sites appears likely to directly produce intraspecies phenotypic diversity, providing substrates for Mendelian selection. Thus, if TE-induced regulatory changes confer adaptive advantages, such changes might catalyze divergence and eventual speciation.

## Conclusions

TEs have broadly influenced CTCF binding in human and mouse. The range of TE types and ages we observed in this study suggest this process has been ongoing throughout mammalian evolution, leading to a large proportion of the CTCF-binding variation seen across mammalian species. Interestingly, while ancient TEs have contributed conserved binding sites, TEs appear to contribute primarily to sites that are variable across species and cells. We found the same association between TEs and variability among CTCF binding sites that function as chromatin loop anchors, noting that nearly 1/3 of variable loops across cells and species can be unambiguously assigned to a TE. We speculate that TE-driven CTCF binding expansions have contributed to regulatory flexibility throughout mammalian evolution. This variability may have served as an evolutionary catalyst, conferring different adaptive advantages to subpopulations of the most-recent common ancestor of human and mouse. This may have allowed them to specialize to different evolutionary niches, eventually leading to speciation. This work will enable important follow-up investigations, including exploration of the regulatory consequences of TE-induced looping variation, which will extend our understanding of the mechanisms effecting such changes. Furthermore, this work advances our understanding of the relationship between TE sequence and functional exaptation, raising important questions about the involvement of CTCF binding and chromatin looping in TE biology.

## Methods

### Overlap of CTCF occupancy between human and mouse

For step 1, CTCF binding sites for human GM12878 and K562, and mouse CH12 and MEL cells were retrieved from the ENCODE repository, using all released datasets (Table S1). Biological and technical replicates were combined using bedtools merge (58) and stored in narrowPeak format. For merged peaks, we assigned the summit location as the centroid of the peak summits for all constituent binding sites. For broadPeak records, the midpoint of the binding site was used as a proxy for the narrowPeak summit. This procedure was repeated in pairs of cell-wise files from the same species in order to determine the union set of CTCF-bound sites in each species.

The next step involved comparison of species-wise sets of CTCF binding sites to determine the extent of overlap between human and mouse CTCF binding landscapes. Prior to cross-species mapping, a unique identifier was assigned to the name column of the input files to facilitate backward comparisons of mapped features across species. CTCF binding peaks were then mapped across species using a modified version of bnMapper (59), with an added option to retain peaks spanning multiple chains by keeping the longest subalignment, following the convention used by the liftOver utility (60). This modified version is freely available at https://github.com/Boyle-Lab/bx-python.

Step 3 involved merging native CTCF binding peak locations and CTCF binding peaks mapped from the other species into union sets representing the locations of all mappable CTCF binding sites across species. Comparisons between the merged narrowPeak files prepared in step 2 were made with bedtools intersect (58) in order to apply labels indicating the specie(s) in which each site was occupied.

The sets resulting from step 3 were then intersected with the locations of all annotated transposable elements from RepeatMasker (46), excluding “Low_complexity”, “Satellite”, “Simple_repeat”, “tRNA”, “rRNA”, “scRNA”, “snRNA”, and “srpRNA” families. Bedtools intersect was used to identify all CTCF sites in which a transposable element was detected within +-50 bp of the ChIP-seq peak summit, and an additional column of labels was added accordingly.

In step 5, procedures from steps 3 and 4 were applied to the species-wise unmapped CTCF ChIP-seq peaks. These were loaded into R data frames along with the union sets produced in step 5, and unique identifiers applied in step 2 were used to identify sites that did not cross-map with bnMapper. These were appended to human- and mouse-referenced union sets to yield complete sets of all known CTCF-bound sites across both genomes. The contribution of TEs to shared and species-specific binding sites was visualized using the VennDiagram R library. Sizes and shapes of individual plot segments were adjusted manually to approximate their proportional contribution to the union datasets (Figure 1A).

The expected numbers of TE-derived human-specific and mouse-specific binding sites were calculated based on overlaps between species-specific CTCF binding sites and randomly selected windows following the size distribution of TE-derived CTCF binding sites in each species. We selected N random background regions from the given genome, where N is the number of species-specific CTCF sites derived from TEs. We then counted the number of overlaps between background regions and species-specific CTCF binding sites. We used the median number of overlaps observed over 1000 random trials as the expected number of TE/CTCF overlaps.

### Enrichment of CTCF binding sites within human and mouse transposable elements

We first sought to identify transposable element families that may have served as a source of novel CTCF binding sites in humans and/or mice. This inquiry extends the findings presented in three key papers, in which enrichments of transcription factor binding sites, including CTCF, were identified in several TE families in humans and mice (27,28,61). Intriguingly, Schmidt et al., the only study to look specifically for enrichment of CTCF binding sites in primate cells, failed to find any significant enrichments in human, despite strong enrichments in mammalian species stretching back to opossum. We wondered whether the reliance of their methods on enrichment of species-specific k-mers at CTCF binding sites influenced their findings. We tested for enrichments using two approaches: binomial tests based on methods used in Bourque et al., and permutation tests based on methods presented in Chuong et al. (61). In both methods, we rely solely on the observed frequency of CTCF binding within each TE type compared to a random expectation to determine enrichments. We performed both analyses on genome-wide sets of CTCF binding sites, and on a more restricted set of CTCF sites located in mouse and human cis-regulatory modules (CRMs) previously reported by us (62).

Merged CTCF ChIP-seq peak files were loaded into a Hadoop Hive database after adding unique id, species, cell, and target columns. The locations of all annotated transposable elements for human and mouse were retrieved from RepeatMasker (46), (Table S1). Data were converted to bed format and all records, excluding “Low_complexity”, “Satellite”, “Simple_repeat”, “tRNA”, “rRNA”, “scRNA”, “snRNA”, and “srpRNA” families. These were annotated with a unique id, species, and the name and distance to the transcription start site of the nearest gene according to bedtools closest (58) and the knownGenes table for hg19 or mm9 genomes (63). These were loaded into the database and intersections with CTCF binding sites were identified with a series of hive queries. Intersections were based on the CTCF ChIP-seq summit location, which was required to fall within the boundaries of a TE annotation. The resulting data were output as a tab-delimited text for further processing in R.

We tested for enrichment of CTCF binding sites within individual TE types using three methods: individual binomial tests using the average genome-wide rate of CTCF binding within TEs as the expected binomial frequency; individual binomial tests using binomial expected frequencies based on CTCF binding within each TE type in permuted data; and enrichment tests based on empirical cumulative density functions computed from the permuted datasets.

For our first set of individual binomial tests, we calculated the genome-wide fraction of TEs containing CTCF binding sites within all four cell types separately and used these as the binomial expected frequencies. Within each cell type, we performed individual binomial tests for every TE type in which CTCF binding was observed and adjusted p-values for multiple testing using the bonferroni method. We applied three criteria for significant enrichment within a TE type: p-value <= 1e-4, at least 25 CTCF-bound TE insertions, and a CTCF binding rate of at least 1% within the given TE type.

For permutation tests, we used a method based on Chuong et al. (61). Starting with the R data frames used in binomial tests, we performed 10,000 random permutations of the CTCF-TE data by shuffling the associated TE types. For each permutation, a Fisher-Yeats shuffle was performed on the name column of the whole-genome repeatMasker annotations using fyshuffle (fgpt R package). In order to maintain the insertion biases of each TE type, shuffling was performed separately within six distance-based bins relative to the nearest transcription start site for each TE insertion. Shuffled names were then applied to the corresponding records for CTCF-bound repeats. The number of times a given TE type was observed among records originally labeled with that family was recorded at each permutation, and resulting counts were used to generate an empirical CDF for each TE family using the built-in ecdf function. Empirical p-values for enrichment of CTCF sites in each TE type were computed by plugging the observed number of CTCF binding events into the corresponding CDF functions, and a Bonferroni multiple testing correction was applied. We applied the same set of criteria used in our binomial tests to assess significance among these results.

### Calculation of motif-word frequencies within enriched TE types

We performed our motif-word frequency enrichment analysis by replicating the procedures presented in (27) and applying them to our own CTCF motif predictions within human and mouse CTCF-bound repeats. Initially, CTCF-bound repeat insertions in the human and mouse genomes were identified by intersecting repeatMasker annotations, excluding “Low_complexity”, “Satellite”, “Simple_repeat”, “tRNA”, “rRNA”, “scRNA”, “snRNA”, and “srpRNA” families, with ChIP-seq peak summit locations using bedtools intersect (58), and fasta sequences were extracted from the hg19 and mm9 genomes with bedtools getfasta (58). FIMO motif prediction was performed using a previously published CTCF position weight matrix (64), using default parameters and a maximum site count of 1,000,000. Predictions were converted to bed format, retaining the sequence of the predicted binding sites as the final bed field. These were read into R data frames and motif-words – distinct 20-mers contributing to the pools of CTCF binding site predictions, were enumerated within human and mouse. Observed bound word-counts were scaled by a normalization factor:

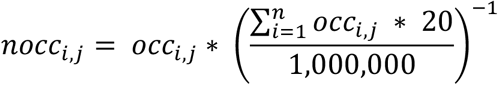

where *i* denotes the word number, *j* denotes the species, and 20 is the length of the CTCF motif in bp. Odds ratios denoting species-specificity were then calculated:

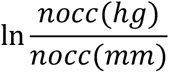

where *nocc(hg)* and *nocc(mm)* are the normalized word counts for human and mouse, respectively. As per Schmidt et al., we considered all motif-words with a normalized occurrence rate of at least 8 and an absolute odds ratio of 2 or greater as species-specific.

We next tested for association of individual human and mouse TE types with species-specific motif-words by comparing their occurrence rates within CTCF-bound TE elements of a given type to their occurrence rate in the rest of the genome. We first isolated CTCF-bound motif-words within the human and mouse genomes by intersecting genome-wide CTCF motif predictions, prepared according to the procedures described above, with 100bp windows surrounding CTCF ChIP-seq peak summit locations in each species using bedtools intersect (58). These were read into an R data frame. For each TE type observed in the CTCF-bound repeat data, individual Fisher’s exact tests were performed, comparing the observed number of species-specific versus shared motif-words in bound repeats compared to background sequences, defined as all CTCF-bound sequences in the given genome excluding those of the given repeat type. P-values were adjusted by applying a Bonferroni correction, and a one-sided p-value threshold of < 10e^-40^ was used to determine significance, as in Schmidt et al.

### Age estimation for CTCF-associated transposable elements

We estimated the ages of TE insertions by dividing the percent divergence from the repeat-wise consensus sequence, as reported by RepeatMasker, by an estimate of the mutation rate per base per year for each species. Although there is some disagreement in the community about the “correct” mutation rates for human and mouse, we chose to use estimates presented by Kumar and Subramian (65). For human, we used the consensus mammalian rate of 2.2*10^-9^ substitutions/base/year, which agrees with the widely-accepted rate presented by the Human Genome Sequencing Consortium (66). For mouse, we used a rate of 2.4*10^-9^, obtained by dividing the human rate by 0.091, to account for the 9.1% faster mutation rate in rodents relative humans reported by Kumar and Subramian (65). This rate is substantially slower than the rate of 4.5*10^-9^ reported by the Mouse Genome Consortium (67), which was used to prepare age estimates in Schmidt et al. (27). However, Kumar and Subramian make a compelling argument that biased substitution patterns can artificially inflate estimated mutation rates (65) and, thus, we opted to use the slower rate. Box plots were then prepared in R using the ggplot2 suite to produce Figure 1E.

### Phylogenetic gain and loss analysis of CTCF binding sites

In order to assign labels denoting evolutionary history to each CTCF binding site present in the human and mouse genomes, we developed a method, mapGL, based on bnMapper (59). MapGL uses chained alignments between a target and query species, and one or more outgroup species to infer whether genomic elements present in the query species are orthologous, gained in the query species, or lost in the target species. Briefly, query elements that map to the target species are labeled as orthologs and printed to output. Non-mapping elements are further mapped to each outgroup species and labels indicating presence/absence of an orthologous element are stored at the corresponding leaf nodes of the phylogenetic tree connecting all species. A maximum parsimony algorithm is then applied to infer whether an orthologous element was present at the root node, corresponding to the most-recent common ancestor (MRCA), of the phylogenetic tree. Elements predicted as present in the MRCA are labeled as losses in the target species, while those predicted to be absent in the MRCA are labeled as gains in the query species. MapGL is freely available at https://github.com/adadiehl/mapGL.

We obtained liftover chains for human (hg19) to mouse (mm9), and for human and mouse to three outgroup species: dog (canFam2), horse (equCab2), and elephant (loxAfr3), from the UCSC Genome Browser download portal (68). We then constructed reciprocal-best alignment chains, representing one-to-one relationships of syntenic blocks between each genome, based on methods described at http://genomewiki.ucsc.edu/index.php/HowTo: Syntenic Net or Reciprocal Best.

We ran mapGL on human and mouse inputs separately, using the reciprocal-best human-to-mouse chain to map human elements, and the reversed human-to-mouse chain to map mouse to human. Relationships between the target and query species and outgroup species are described by the Newick tree: (((hg19,mm9),(canFam2,equCab2)),loxAfr3). mapGL.py was invoked with the “–input_format narrowPeak” option, to include the mapped location of narrowPeak summits in output whenever possible.

We first intersected 50bp windows surrounding each human and mouse ChIP-seq peak summit with repeatMasker repeats, as described previously. Phylogenetic labels were then applied with mapGL, and the results were annotated with additional data from the original repeatMasker files. These were further analyzed in R. Specifically, the contribution of each phylogenetic class to each CTCF-enriched repeat type was assessed by plotting the fraction of elements within each type assigned as orthologous, gained, or lost on a given branch using the ggplot2 R package.

### Motif scores within CTCF-enriched ancestral repeats

We first computed log-odds scores for matches to a previously-published CTCF position weight matrix (PWM) (64) at every position in the human and mouse genomes using a custom Perl script (score_motifs.pl, available at …). These were stored in wig format and converted to bigWig files using the wigToBigWig utility (60). We made use of the rtracklayer package to retrieve motif scores from these bigWig files for TE instances annotated as orthologous across human and mouse. For each of these regions, scores were retrieved for a 50bp window centered around the summit of the embedded CTCF ChIP-seq peak. This was repeated for the orthologous location in human or mouse, and the maximum scores were stored for both species. To obtain the CTCF motif score distribution within consensus elements for human-only TEs, we first extracted the consensus sequence, in fasta format, for each TE type from Repbase version 23.09 (48). Each consensus sequence was scored using score_motifs.pl and the same CTCF PWM used to generate the bigWig files, and the maximum scores from each were retained. Score distributions were visualized using the ggplot2 R package. We assessed the significance of the observed difference between human and mouse score distributions and mouse using a Wilcoxon signed-rank test, and between consensus, human, and mouse using Wilcoxon rank-sum tests.

### MANGO analysis of RAD21 ChIA-pet data

We retrieved ChIA-pet data for human GM12878 and K562 from the ENCODE download portal in fastq format (Table S1). After careful evaluation of FastQC (69) results on each input file, we elected to proceed without adapter trimming. Paired reads in fastq format were aligned separately with BWA mem (70) with default parameters. Mapping quality was evaluated with SAMtools flagstat (71) found to be >= 94% for all but two files. Samtools view was then used to filter out unmapped and secondary reads (SAM flags 4 and 256), and those with quality scores less than 30. Filtered reads were then sorted by X and Y coordinates and a custom script was used to assemble paired reads into bedpe format. Bedpe files for all biological and technical replicates were then concatenated and processed with the MANGO pipeline, starting with stage 3.

### Contribution of TE-borne CTCF binding sites to ChIA-pet and Hi-C loop anchors

RAD21 ChIA-pet loops for human GM12878 and K562 cells, and Hi-C loops for the same human cells and mouse CH12 cells, were first filtered to include only loops containing a CTCF ChIP-seq peak at both anchors. If multiple CTCTF ChIP-seq peaks overlapped a loop anchor, we kept only the peak with the strongest signal value. Loop anchor coordinates were then trimmed to the boundaries of their respective overlapping ChIP-seq peaks. The location of the overlapping ChIP-seq peak summit was included in the record as an additional field. The trimmed and filtered loop loci were then read into a data table in Hadoop hive database table. We then intersected these loops with the CTCF-TE associations prepared in our analysis of CTCF-enriched TEs by comparing the CTCF peak summit locations.

ChIA-pet and Hi-C loops were separately intersected with CTCF motif predictions. CTCF motif predictions were prepared in-house using a custom script. We used a previously-published CTCF position weight matrix (64) and calculated simple log-odds scores relative to the equilibrium nucleotide frequencies for each base for overlapping, 20bp windows spanning the human and mouse genomes. In order to maximize the fraction of CTCF-bound sequences to which motifs could be assigned, we retained all predictions with log-odds scores exceeding 0, although nearly 70% of CTCF-bound sites contained a motif with a log-odds score >= 10. We associated CTCF motifs with RAD21 ChIA-pet and Hi-C loops by extending a 50bp window surrounding the CTCF ChIP-seq peak summit. We then used an R script to assemble a data frame for each cell type, containing a row for each RAD21 ChIA-pet or Hi-C loop, and columns indicating TE presence and CTCF motif presence and orientation in the right and left loop anchors. These data frames were used to collect counts presented in Figure 2 and Table S6.

### Contribution of transposable elements to conserved and species-specific chromatin loops

In order to categorize loops based on sharing between cells, we devised a simple classification scheme based on physical overlap of loop anchors. Loops in the query species were classified by looking for overlaps between their left and right anchors and loop anchors in a set of target loops from another cell. If the left and right anchors both mapped to anchors from the same loop in the target set, a loop was assigned to class C – fully conserved. Class B2 designated loops where both query anchors overlapped target anchors, but target anchors were from different loops. Class B1 designated loops where only one query anchor overlaps a target anchor, and B0 designates loops where neither query anchor overlaps a target anchor. To accommodate cross-species comparisons, we added classes N0, N1A, and N1B, which represent anchors where one or both loop anchors are present in non-orthologous sequence. N0 denotes loops where both anchors are non-orthologous to the target species. N1A denotes sequences where one query anchor is both orthologous and overlaps a target loop anchor. N1B denotes loops where one query anchor is orthologous but does not overlap a target loop anchor. Loops from all three cell types for which we have loop data (GM12878, K562, and CH12) were assigned to these classes using another adaptation of bnMapper, which we call mapLoopLoci. This tool is available from our github repository: https://github.com/adadiehl/mapLoopLoci. We assigned conservation classes between loops for all pairwise combinations of cells and combined the results into an R data frame. We then intersected these data with the results from our previous analysis of TE-loop intersections based on previously assigned loop ID numbers and counted the fractions of loops in each conservation class contributed by TEs. In the case of species-specific and cell-specific loops, we required that the TE insertion overlap the loop anchor(s) unique to the query cell in order to count toward the total number of TE-derived loops.

### Analysis of correlation between loop strength, loop conservation, and transposable element content

To determine whether any correlation exists between loop strength, loop conservation, and TE content, we compiled PET counts for RAD21 ChIA-pet loops in human GM12878 and K562 cells, and observed Hi-C contact counts for mouse CH12 cells, in each of the seven conservation classes defined in “Contribution of transposable elements to conserved and species-specific chromatin loops.” Box plots for TE-derived and non-TE-derived fractions for each set of scores were produced using the ggplot2 R package in order to visualize any score trends. This process was repeated for all pairwise combinations of cells. Wilcoxon rank-sum tests were performed to determine the significance of any trends toward higher or lower scores for all pairs of conservation classes within each comparison, and between TE-derived and non-TE-derived loops within each conservation class across all comparisons. Resulting p-values were then plotted as heatmaps with the ggplot2 R package.

### Ages of TE elements in each loop conservation class

To determine if any trends were evident between conservation classes and the estimated ages of TEs contributing to each class, we estimated the ages for all TE insertions within each conservation class in mouse-to-human and human-to-mouse comparisons according to procedures reported in “<u>Age estimation for CTCF-associated transposable elements</u>”. Score distributions for each conservation class were rendered as box-plots and visually compared to identify any notable trends. In order to test for a significant linear correlation between conservation classes and estimated TE ages, we assigned numeric values to each conservation class and fitted a linear model relating conservation class and estimated TE age with the “lm” function in R. Goodness-of-fit and significance of the observed linear trend were determined by applying the summary function to the fitted lm model. We further tested for significant differences in estimated TE age distributions between pairs of conservation classes and TE age in mouse-human comparisons using individual Wilcoxon rank-sum tests. The p-values of these tests were compiled into matrix form and visualized with ggplot2 to produce the heatmap presented in Figure 3E.

### Relative contributions of CTCF-Enriched TE types to loop conservation classes

To determine if there is a relationship between loop conservation and CTCF-enriched TE types, we labelled each TE insertion in the loop conservation dataset as mouse-enriched, human-enriched, shared, or non-enriched. The contribution of each enrichment category to loops in each conservation class was visualized by iteratively applying the “table” function in R and plotting the resulting table of counts as stacked bar graphs with ggplot2. For the mouse-human comparison (Figure 3F), the observed counts for human-enriched and shared TE types were combined for clarity. This only affected the “C” and “B2” conservation classes.

### Analysis of correlation between loop conservation classes and evolutionary conservation

We first retrieved phastCons 46-way placental mammal conservation scores in wig format from the UCSC download portal (63). These were subsequently converted to bigwig format with the wigToBigWig utility (60). We used an R function, making use of the rtracklayer Bioconductor package, to retrieve phastCons scores for 500bp windows surrounding the annotated CTCF peak summit location within all TE-borne loop anchors and the resulting matrix of scores was summarized with the colMeans function. This process was applied to all loops in each conservation class and mean score vectors were stored in a data frame and plotted as line graphs with ggplot2.

### Overlap with topologically-associating domains

We obtained published predicted locations of topologically-associating domains (TADs) for CH12, GM12878, and K562 cells from the GEO repository for (3) (Table 1). We converted the primary data from the Arrowhead format to a BED format where the start and end coordinates correspond to the X and Y coordinates defining the TAD boundaries within the Hi-C contact matrix and read into R for further processing. In order to identify loops in our dataset that correspond to known TADs, we first found the intersection with our loop dataset using the findOverlaps method from the GenomicRanges R library, with default options. This identified which loops overlapped known TADs but did not indicate which loops correspond to entire TAD regions. To do so, we defined upstream and downstream boundary regions for each TAD by extending a window +-10kb around their x and y coordinates. We then looked for loops where the upstream and downstream anchors overlapped the upstream and downstream boundaries of a single TAD from the same cell type.

**Table 1:**
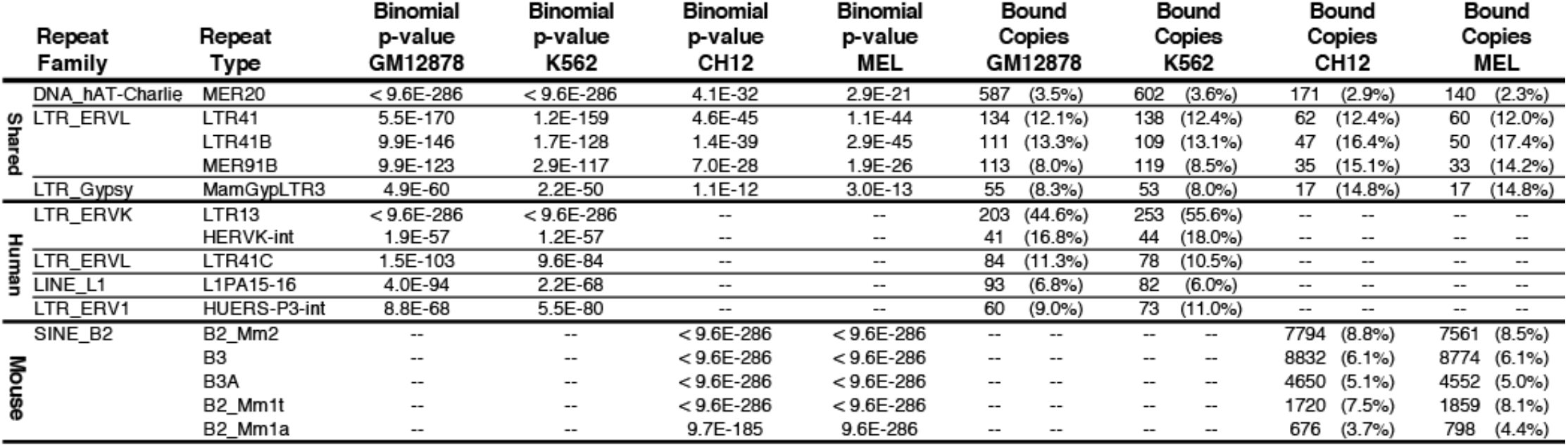
Top-five transposable element types enriched for CTCF binding in shared, human-only, and mouse-only classes.

### Epigenetic properties of conserved and species-specific chromatin loops

We obtained ChIP-seq data for CH12, GM12878, and K562 cells in bigWig format from the ENCODE download portal (Table 1). We used the rtracklayer package to retrieve histone marks signals at each location in 20kbp windows surrounding the annotated CTCF peak summits within loop anchors and stored them in data frame. For each histone mark and cell line, we compared the average signals in left and right portions of each 20kbp window region and flip the direction if the right half has higher signal than left. Mean signal at each location was calculated using the colMeans function and then divided by mean signals at randomly selected 20kbp windows across the genome calculated following the same method. This process was applied to TE-derived and non-TE-derived loops anchors separately and processed score vectors were plotted as line graphs for each cell type using ggplot2. Then for human GM12878 and K562 cells, we filtered regions that are loops anchors in only one of the two cell types and followed the same steps to plotted line graphs for these cell type-specific loop anchors in each cell line.

## Declarations

### Ethics approval and consent to participate

Not applicable

### Consent for publication

Not applicable

### Availability of data and material

The datasets supporting the conclusions of this article are included within the article and its additional files, listed in Table S1. Software and scripts used for data processing and Figure preparation are available in the github repository, https://github.com/Boyle-Lab/TE-Driven-CTCF-Loop-Evol.

### Authors’ contributions

AGD and APB conceived and planned the study. All experiments and analyses were performed by AGD, with the exception of the analysis of epigenetic properties of conserved and species-specific chromatin loops, which was performed by NO. AGD prepared the manuscript with input from all authors. APB supervised the experiments, analysis, and data interpretation. All authors read and approved of the final manuscript.

## Acknowledgements

We would like to thank members of the Boyle lab and Dr. John Vincent Moran for critical reading and suggestions on the manuscript and analyses.

## Additional Files

**Additional File 1: Supplementary Figures**. Supplementary_Figures.pdf. Figures S1-S9.

**Additional File 2: Table S1**. Table-S1_Datasets.xlsx. Datasets used in this analysis.

**Additional File 3: Table S2**. Table-S2_ctcf_te_enrichments.binomial.xlsx. Results for binomial tests of CTCF enrichment in human and mouse transposable elements.

**Additional File 4: Table S3**. Table-S3_ctcf_te_enrichments.permutation.xlsx. Results for permutation tests of CTCF enrichment in human and mouse transposable elements.

**Additional File 5: Table S4**. Table-S4_species-specific-word-enriched-TEs.xlsx. Results for word-enrichment tests of CTCF motif enrichment in human and mouse transposable elements.

**Additional File 6: Table S5**. Table-S5_top-enriched-words.binomial.xlsx. Enriched motif-words in CTCF motif-word enrichment tests in human and mouse transposable elements.

**Additional File 7: Table S6**. Table-S6_Loop-TE-Motif-Intersections.xlsx. Accounting of TE-loop intersections in human and mouse cells.

**Additional File 8: Table S7**. Table-S7_Loop-conservation.xlsx. Loop conservation statistics for TE-derived and non-TE-derived loops.

