## Supplemental Figures 1-9 for "Transposable elements strongly contribute to cell-specific and species-specific looping diversity in mammalian genomes"

**Table of Contents**

**Figure S1:** CTCF enrichments are robust to different statistical testing procedures and stable across cell types.
**Figure S2:** Differences in functional annotation associations do not explain CTCF-binding enrichments in human and mouse TE types.
**Figure S3:** TE-loop associations are stable across species and cell lines.
**Figure S4:** TE-derived vs not TE-derived loop anchors comparison.
**Figure S5:** Conservation class assignment algorithm.
**Figure S6:** Overlaps between loops in the present dataset and known TADs and TAD boundaries.
**Figure S7:** All conservation classes of TE-derived loops show evidence of natural selection.
**Figure S8:** Associations between transposable elements and conservation classes are stable across cell lines.
**Figure S9:** Comparison of histone modifications for non-conserved loops.

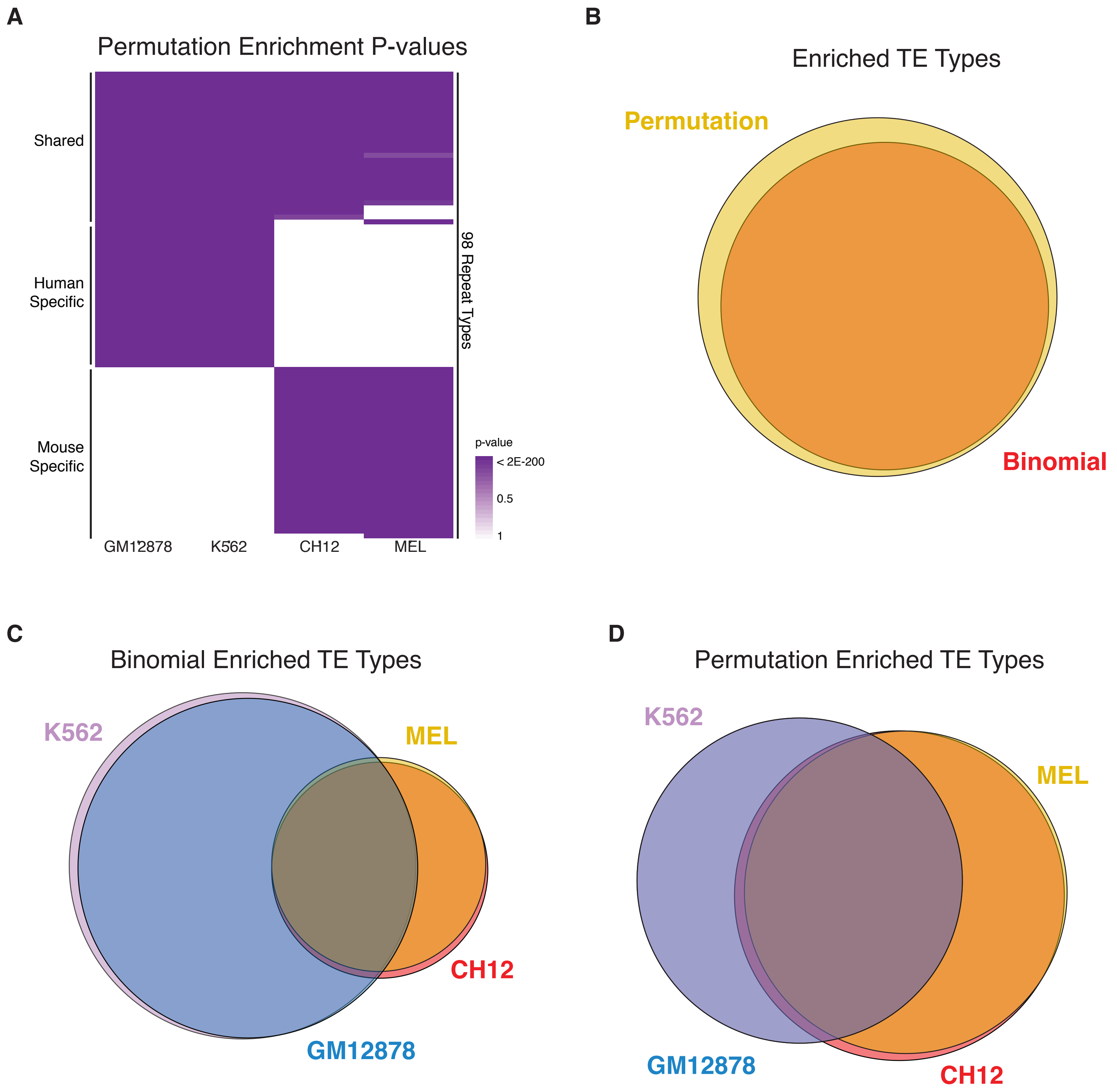


**E**

| | Repeat Family | Repeat Type | p-value GM12878 | p-value K562 | p-value CH12 | p-value MEL | Bound Copies G | Bound Copies K | Bound Copies C | Bound Copies M | % Bound Copies G | % Bound Copies K | % Bound Copies C | % Bound Copies M |
|---|---|---|---|---|---|---|---|---|---|---|---|---|---|---|
| Shared | DNA_hAT-Charlie | MER20 | < 1.6E-284 | < 1.6E-284 | < 1.6E-284 | < 1.6E-284 | 587 | 602 | 171 | 140 | 3.5% | 3.6% | 2.9% | 2.3% |
| | LTR_ERVL | LTR41 | < 1.6E-284 | < 1.6E-284 | 1.6E-214 | 4.4E-207 | 134 | 138 | 62 | 60 | 12.1% | 12.4% | 12.4% | 12.0% |
| | | LTR41B | < 1.6E-284 | < 1.6E-284 | 4.0E-171 | 2.2E-182 | 111 | 109 | 47 | 50 | 13.3% | 13.1% | 16.4% | 17.4% |
| | | LTR50 | < 1.6E-284 | < 1.6E-284 | 1.5E-79 | 2.8E-61 | 185 | 199 | 24 | 19 | 7.3% | 7.8% | 4.4% | 3.5% |
| | DNA_hAT-Tip100 | MER91B | < 1.6E-284 | < 1.6E-284 | 1.1E-131 | 1.0E-124 | 113 | 119 | 35 | 33 | 8.0% | 8.5% | 15.1% | 14.2% |
| Human | LTR_ERVK | LTR13 | < 1.6E-284 | < 1.6E-284 | -- | -- | 203 | 253 | -- | -- | 44.6% | 55.6% | -- | -- |
| | LTR_ERVL | LTR41C | 1.7E-283 | 2.6E-260 | -- | -- | 84 | 78 | -- | -- | 11.3% | 10.5% | -- | -- |
| | LINE_L1 | L1PA17 | 1.1E-170 | 8.5E-156 | -- | -- | 68 | 62 | -- | -- | 1.4% | 1.3% | -- | -- |
| | | L1PA15-16 | 1.6E-284 | 1.6E-251 | -- | -- | 93 | 82 | -- | -- | 6.8% | 6.0% | -- | -- |
| | LTR_ERV1 | HUERS-P3-int | 2.2E-204 | 3.8E-251 | -- | -- | 60 | 73 | -- | -- | 9.0% | 11.0% | -- | -- |
| Mouse | SINE_B2 | B2_Mm1a | -- | -- | < 1.6E-284 | < 1.6E-284 | -- | -- | 676 | 798 | -- | -- | 3.7% | 4.4% |
| | | B2_Mm1t | -- | -- | < 1.6E-284 | < 1.6E-284 | -- | -- | 1720 | 1859 | -- | -- | 7.5% | 8.1% |
| | | B2_Mm2 | -- | -- | < 1.6E-284 | < 1.6E-284 | -- | -- | 7794 | 7561 | -- | -- | 8.8% | 8.5% |
| | | B3 | -- | -- | < 1.6E-284 | < 1.6E-284 | -- | -- | 8832 | 8774 | -- | -- | 6.1% | 6.1% |
| | | B3A | -- | -- | < 1.6E-284 | < 1.6E-284 | -- | -- | 4650 | 4552 | -- | -- | 5.1% | 5.0% |

**Figure S1: CTCF enrichments are robust to different statistical testing procedures and stable across cell types.** **A)** A permutation-based testing approach was used to retest for CTCF binding enrichments. 98 significant results were recovered. **B)** Permutation tests recovered 100% of TE types found to be enriched in binomial tests. **C-D)** The majority of enriched TE types were detectable across all cell types tested in each species in both binomial and permutation tests. **E)** Top five shared, human-only, and mouse-only enriched TE types for permutation tests.

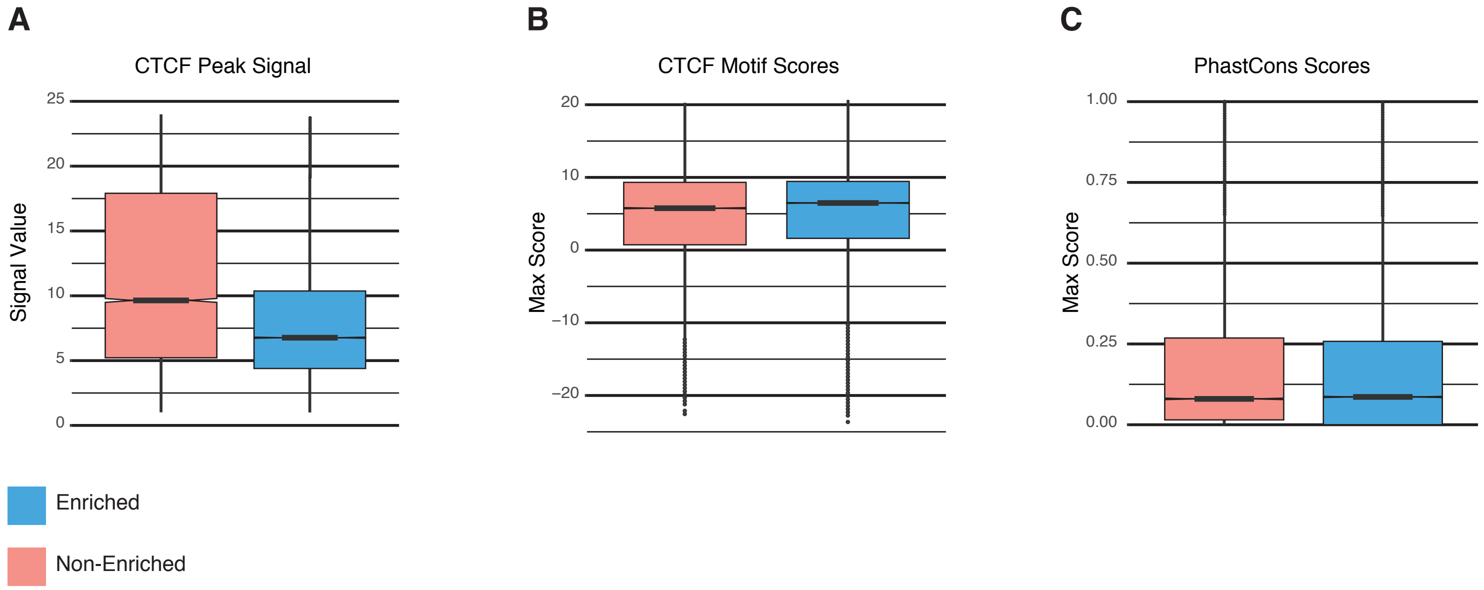


**Figure S2: Differences in functional annotation assocations do not appear to explain CTCF-binding enrichments in human and mouse TE types.** Box plots illustrate score distributions for three functional annotations associated with instances of CTCF-enriched and non-enriched TE types. We wanted to see if enriched TE types displayed any systematic differences in functional properties that might explain their enrichment for CTCF binding. We considered three different functional annotations that might explain differential CTCF enrichment: **A)** CTCF ChIP-seq peak signal intensity. If CTCF binds more strongly to sites within enriched TE types, it might explain why they appear in enrichment tests. Surprisingly, these scores are actually systematically lower in CTCF-enriched TE instances compared to non-enriched TE types. **B)** We see no systematic differences in CTCF motif scores within enriched and non-enriched TE instances. This suggests that extant copies of both classes of TEs may be equally capable of binding CTCF, and fails to explain differential enrichments. **C)** PhastCons conservation scores show no signficant differences in purifying selection between CTCF sites within enriched and non-enriched TEs, suggesting that both classes are equally likely to experience functional constraint.

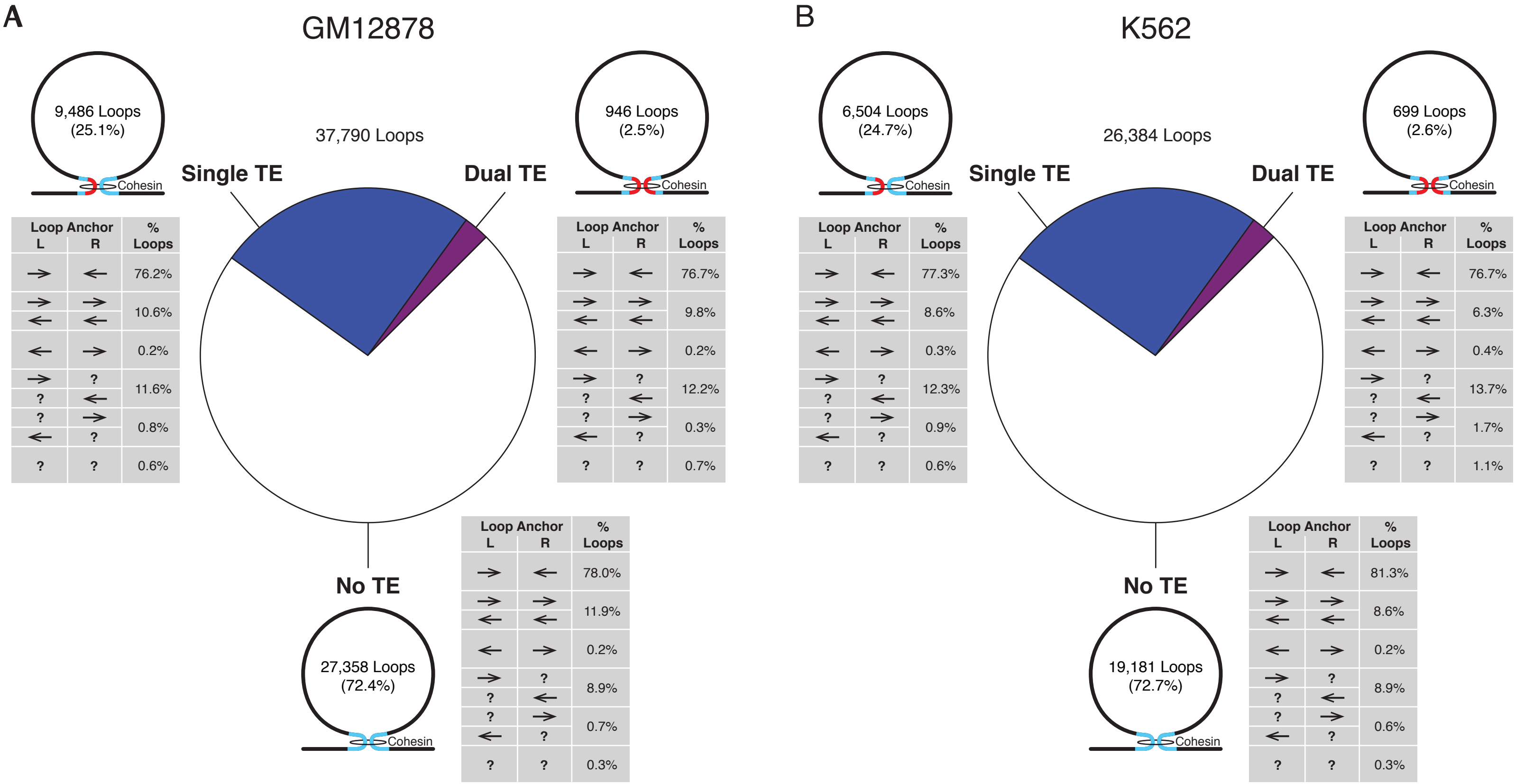


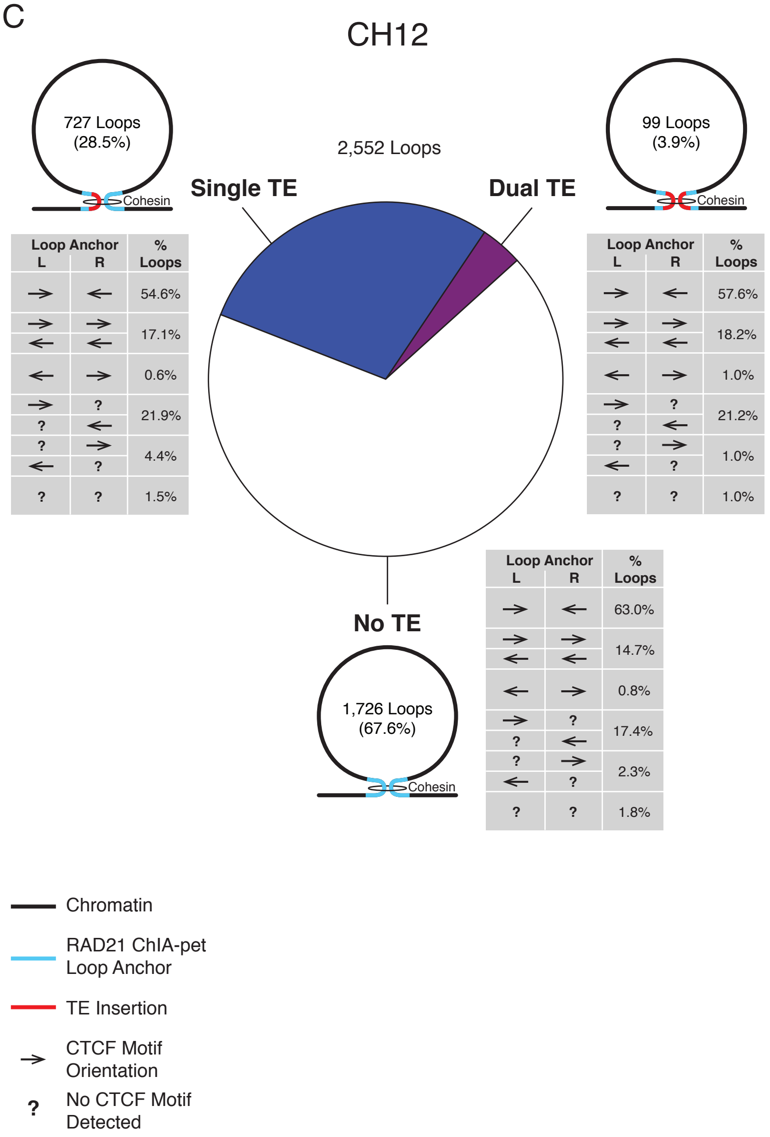


**Figure S3: TE-loop associations are stable across species and cell lines.** Fractions of TE-derived human RAD21-mediated chromatin loops and mouse Hi-C loops across three cell lines. Pie charts illustrate proportions of loops constructed using one, two, or no TE-derived loop anchors. Associated numbers for each of these three classes are given within the corresponding loop diagrams, and the fraction of those loops displaying every possible CTCF motif layout are given in the associated tables. Observations show very little variation across all three cell types: **A)** Human GM12878 cells **B)** Human K562 cells **C)** Mouse CH12 cells.

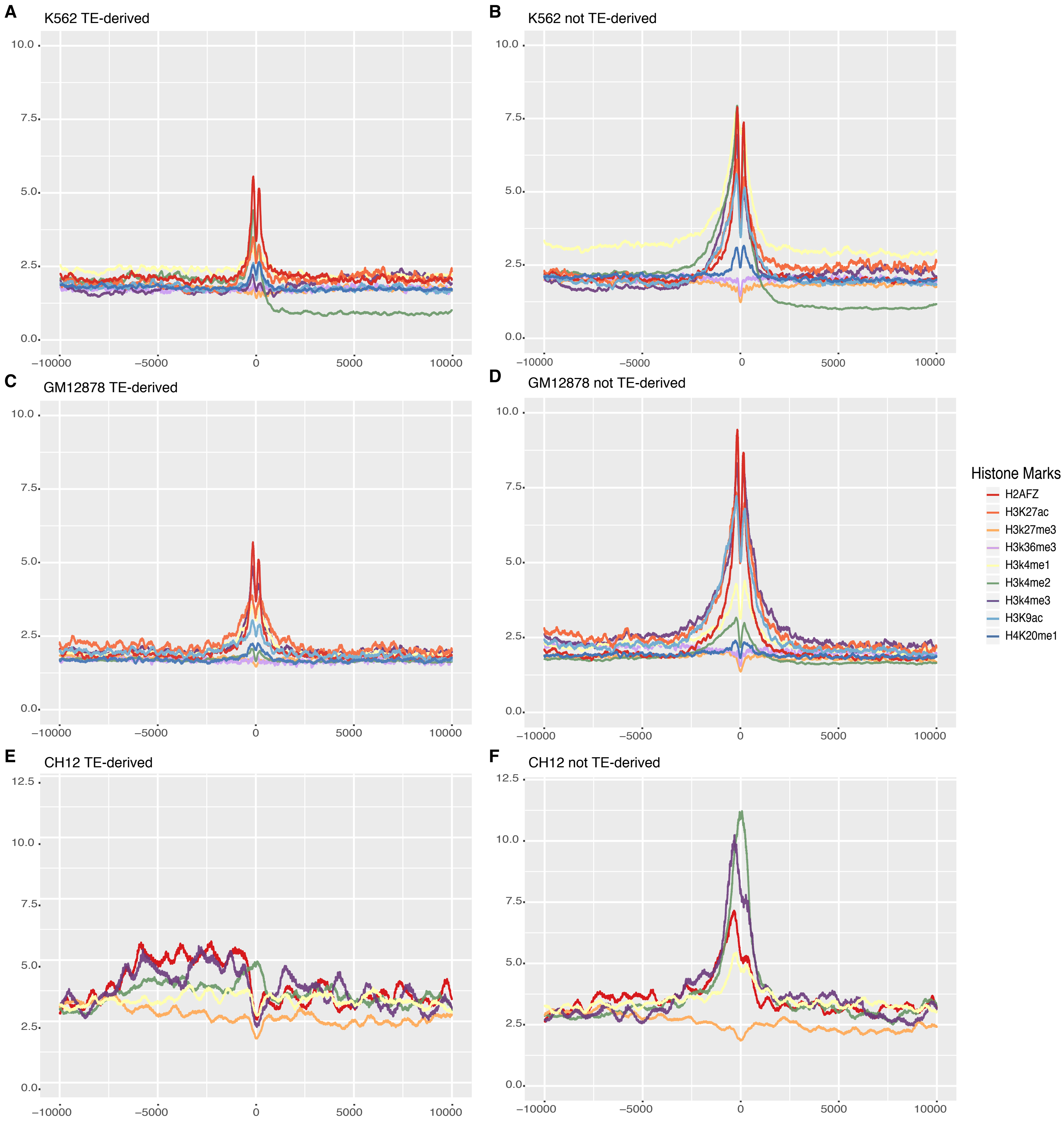


**Figure S4:** TE-derived vs not TE-derived loop anchors comparison. Average histone modification signals are centered at CTCF peak summits in loop anchors that are (a, c, e) TE-derived or (b, d, f) not TE-derived in K562, GM12878, and CH12, normalized by signals at randomly chosen sites in corresponding cell types. e) TE-derived CH12 loops. Although histone mark enrichments around TE-derived loop anchors in CH12 are of similar magnitude to those in the human cell types, they do not form a distinct peak. We believe this is likely an artifact of decreased read mappability within mouse repetitive sequences.

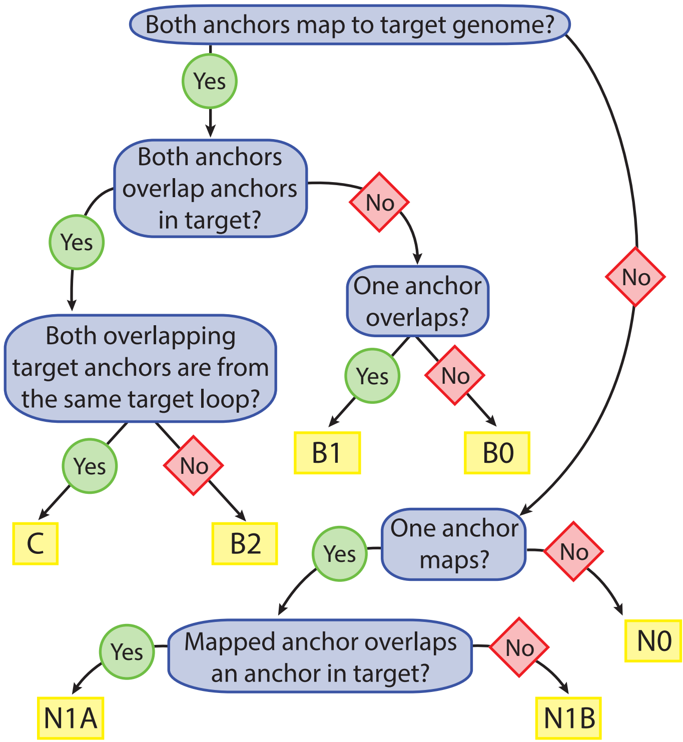


**Figure S5: Conservation class assignment algorithm.** The process of assigning loops to conservation classes can be described as a decision tree, in which anchor mappings between query and target genomes are considered first, after which query anchors that do map to the target genome are checked for overlap(s) with loop anchors detected in the target dataset.

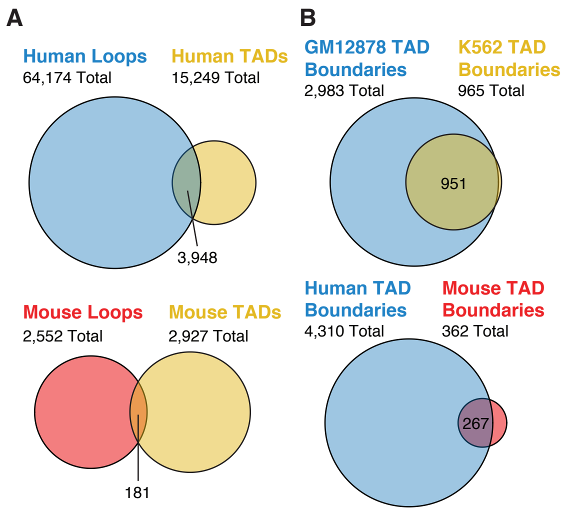


**Figure S6: Overlaps between loops in the present dataset and known TADs and TAD boundaries.** We aligned chromatin loops with previously annotated TADs {Rao:2014eo}, requiring that both loop anchors map to a coherent set of TAD boundaries to call a match. **(A)** Our dataset captured 26% of human TADs and 6% of mouse TADs in the same cell types when strict anchor matching was enforced. **(B)** Upon closer examination, we noted that conservation metrics reported in previous studies were based only on overlap between individual TAD boundaries, not the entire interval spanned by the TADs being compared {Dixon:2012fb}{VietriRudan:2015gl}. When we compared TAD boundary overlap in an analogous manner, our dataset captured 48% of mouse TADs and 55% of human TADs, and the interspecies and intercell overlaps we observed were consistent with previous reports {Dixon:2012fb}{VietriRudan:2015gl}.

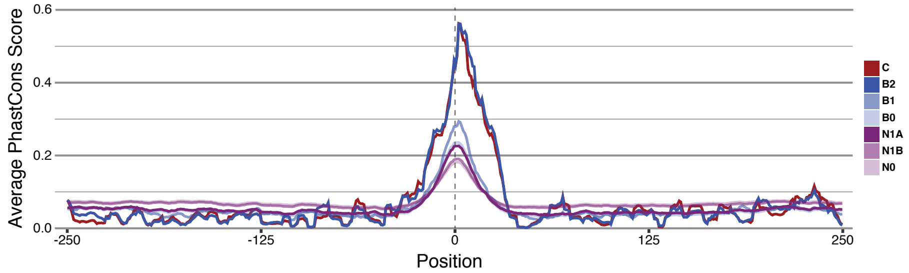


**Figure S7: All conservation classes of TE-derived loops show evidence of natural selection.** PhastCons scores in a 500bp window surrounding the CTCF peak summit embedded in TE-derived loop anchors are plotted. The strength of conservation at the CTCF binding site appears to be correlated with the degree of loop conservation, following the same pattern observed for non-TE loops.

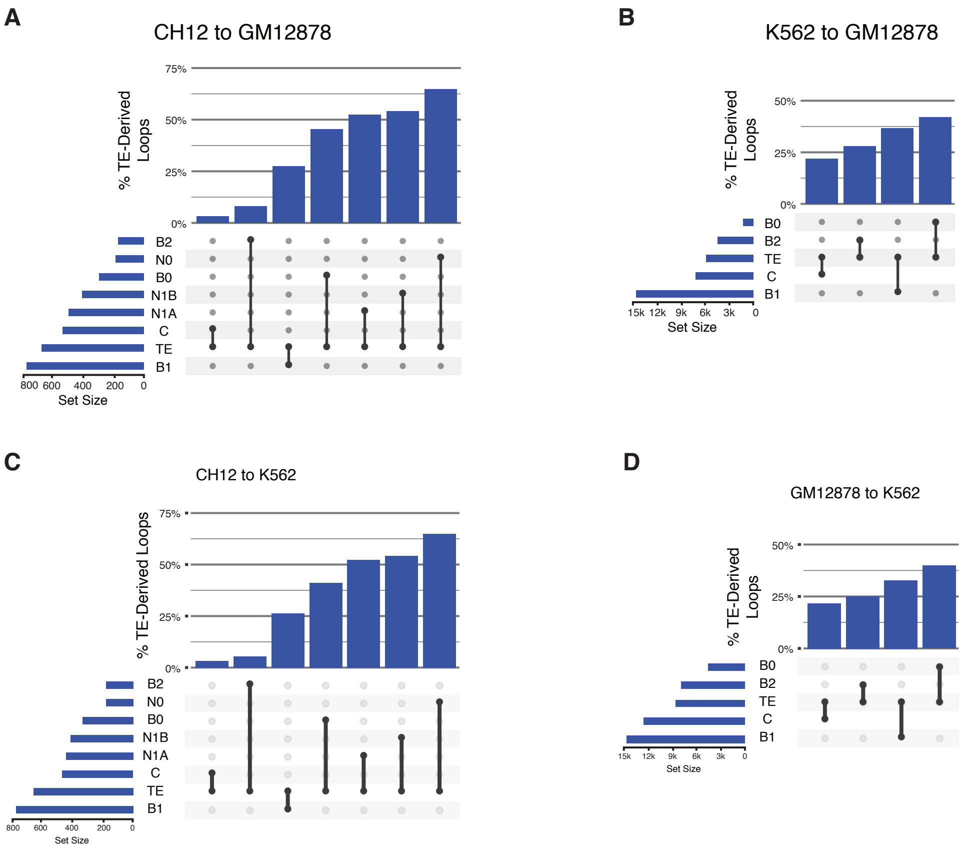


**Figure S8: Associations between transposable elements and conservation classes are stable across cell lines.** The fraction of loops in each conservation class derived from TE insertions is shown for pairwise comparisons between mouse and human cells, and between different human cell types. All comparisons show a trend toward greater TE-derived contributions as conservation decreases. **A)** Mouse CH12 cells compared to human GM12878 cells. **B)** Human K562 cells compared to human GM12878 cells. **C)** Mouse CH12 cells compared to human K562 cells. **D)** Human GM12878 cells compared to human K562 cells.

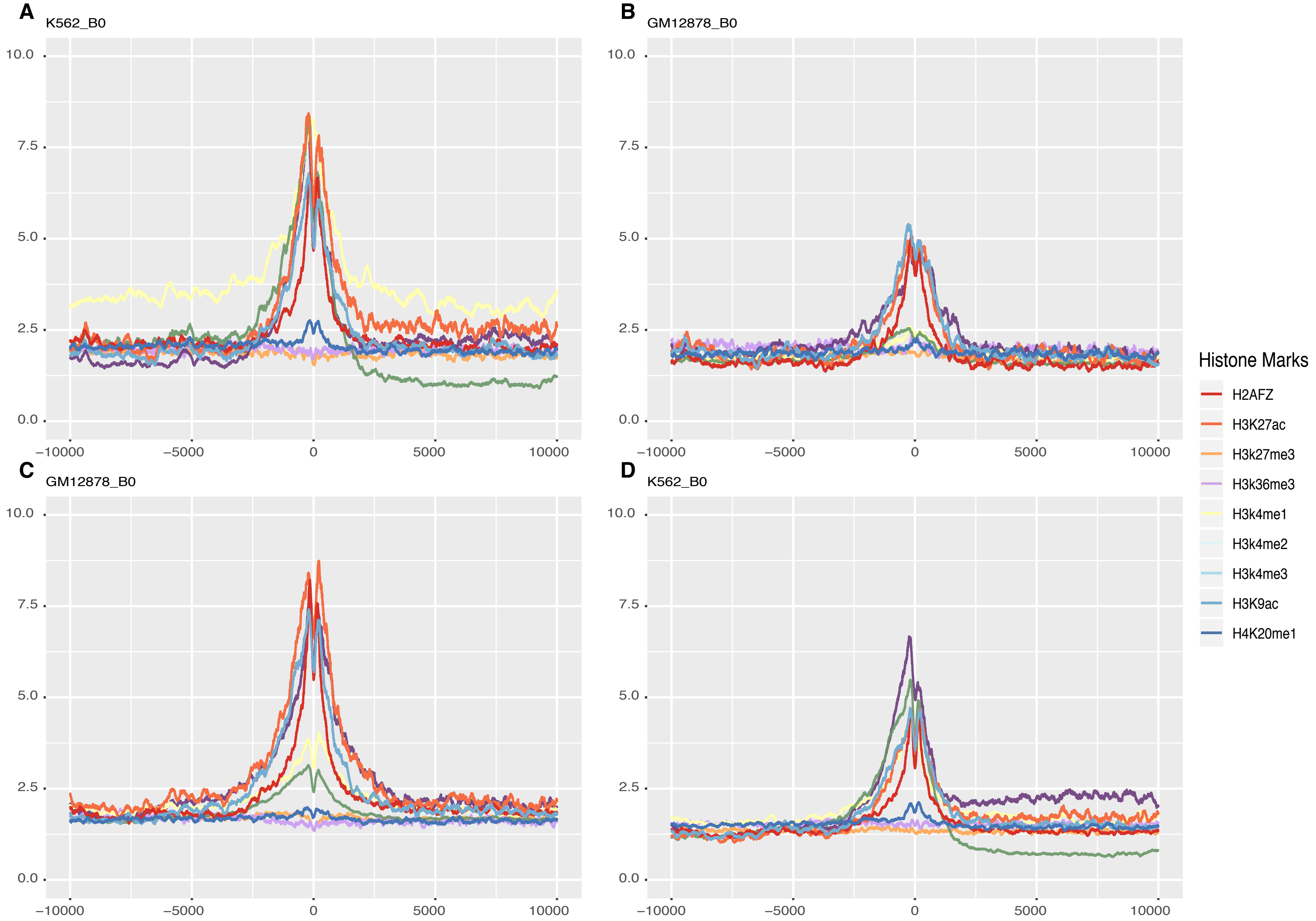


**Figure S9:** Comparison of histone modifications for non-conserved loops. Average histone modification signals are centered at CTCF peak summits in non-conserved loop anchors of the B0 class that are TE-derived in cross-cesl comparisons between human K562 and GM12878 cells. Signals are normalized against randomly chosen sites in corresponding cell types. A) Histone modification scores for K562 loops not found in GM12878. B) Histone modification scores for GM12878 at loci for loop anchors depicted in (A). C) Histone modification scores for GM12878 loops not found in GM12878. D) Histone modification scores for K562 at loci for loop anchors depicted in (C).
